# To join or not to join: handling biological replicates in long-read RNA sequencing data

**DOI:** 10.64898/2025.12.09.693269

**Authors:** Fabian Jetzinger, Alejandro Paniagua, Stanley Cormack, Jorge Mestre-Tomás, José Manuel Morante-Redolat, Isabel Fariñas, Luis Ferrández, Stefan Götz, Carolina Monzó, Ana Conesa

**Affiliations:** BioBam Bioinformatics S.L., Valencia, Spain; Institute for Integrative Systems Biology, Spanish National Research Council, Paterna, Valencia, Spain; Department of Computer Science, University of Valencia, Spain; Department of Life Sciences, Imperial College London, London, United Kingdom; Department of Cellular and Functional Biology, Institute for Biotechnology and Biomedicine (Biotecmed), University of Valencia, Spain; Network for Biomedical Research Center on Neurodegenerative Diseases (CIBERNED), Spain

## Abstract

Long-read RNA sequencing (lrRNA-seq) has revolutionized transcriptomics facilitating the study of alternative splicing and resulting in identification of thousands of novel transcripts. While isoform identification has received significant attention, the handling of biologically replicated lrRNA-seq datasets remains less explored. However, how multiple samples are combined in a lrRNA-seq study may strongly impact transcript identification. This study defines and evaluates two strategies for obtaining consensus transcriptomes from multi-sample lrRNA-seq data: “Join & Call”, where reads from all samples are combined before transcript identification, and “Call & Join”, where transcript identification is performed on individual samples before combining the resulting annotations. We applied these strategies to a highly replicated dataset of mouse brain and kidney tissues, using both PacBio and ONT technologies, across six widely used transcript reconstruction tools. Our results indicate that the optimal strategy depends on the chosen computational tool and research objective. We found that Join & Call is generally more suitable for discovering rarely occurring, novel isoforms, as pooling evidence increases confidence in calling lowly-expressed transcripts. Conversely, Call & Join is computationally more efficient and often preferable for highly replicated datasets when the investigation of rare novel transcripts is not the primary objective. Our findings provide a conceptual and practical framework for multi-sample transcriptome reconstruction, guiding best practices in the context of increasingly large-scale lrRNA-seq studies.

## Introduction

Long-read RNA sequencing (lrRNA-seq) has revolutionized transcriptomic research by enabling the sequencing of full-length RNA transcript molecules, allowing researchers to study alternative splicing and diversity of RNA isoforms in greater detail than before^1^. A key outcome of lrRNA-seq studies is the recurrent identification of *thousands* of previously unannotated isoforms even in organisms whose transcriptomes were thought to be comprehensively characterized^2,3^. However, identifying and accurately reconstructing transcript models from long RNA reads is challenging due to the difficulty of separating technical and biological errors from real biological variation^4^. To address this issue, a wide range of computational tools have been developed, utilizing diverse algorithmic approaches – including reference-based, reference-free and annotation-driven transcript model inference^5–10^. Benchmarking efforts by the LRGASP^11^ and SG-NEx^12^ consortiums, among other initiatives^13–15^, have provided the community with valuable guidance on tool and methodology best practices for different biological applications in single-sample analyses. However, in late 2024, both Pacific Biosciences (PacBio) and Oxford Nanopore Technologies (ONT) –the leading providers of lrRNA-seq technologies– significantly increased sequencing accuracy (>99.20%^16,17^) and sequencing throughput (∼100-140 million high-quality cDNA reads per run). With the associated reduction in costs, lrRNAseq datasets now commonly include an increasing number of biological replicates. This shift has made the challenge of transcript model reconstruction yet more complex, as computational tools have implemented different strategies to address how they integrate information across multi-sample datasets^7,18–23^.

Although multi-sample analyses are now becoming routine, the strategies used to handle them have not been formally defined or tested. We address this gap by introducing two paradigms that capture the implicit choices made by current approaches (**Figure 1a**). In a “Join & Call” (J&C) strategy, first reads from all samples are combined and then transcript identification is performed, and eventually, transcript expression levels at each individual sample are quantified against this jointly called transcriptome in a secondary step. This strategy maximized transcript detection power, but is computationally intensive and may potentially mask sample-specific transcripts^21–23^. In contrast, in a “Call & Join” (C&J) strategy, transcript identification is first performed on each individual sample. The resulting transcriptome annotations for each sample are then combined into a single transcriptome. This strategy is more scalable and computationally efficient but may lack sensitivity to lowly expressed transcripts^7,18–20^. While these paradigms encompass the current practices, their relative strengths and weaknesses have not been systematically examined. Moreover, it is not clear if different transcript reconstruction tools may behave comparatively similarly on each strategy.

**Figure 1:**
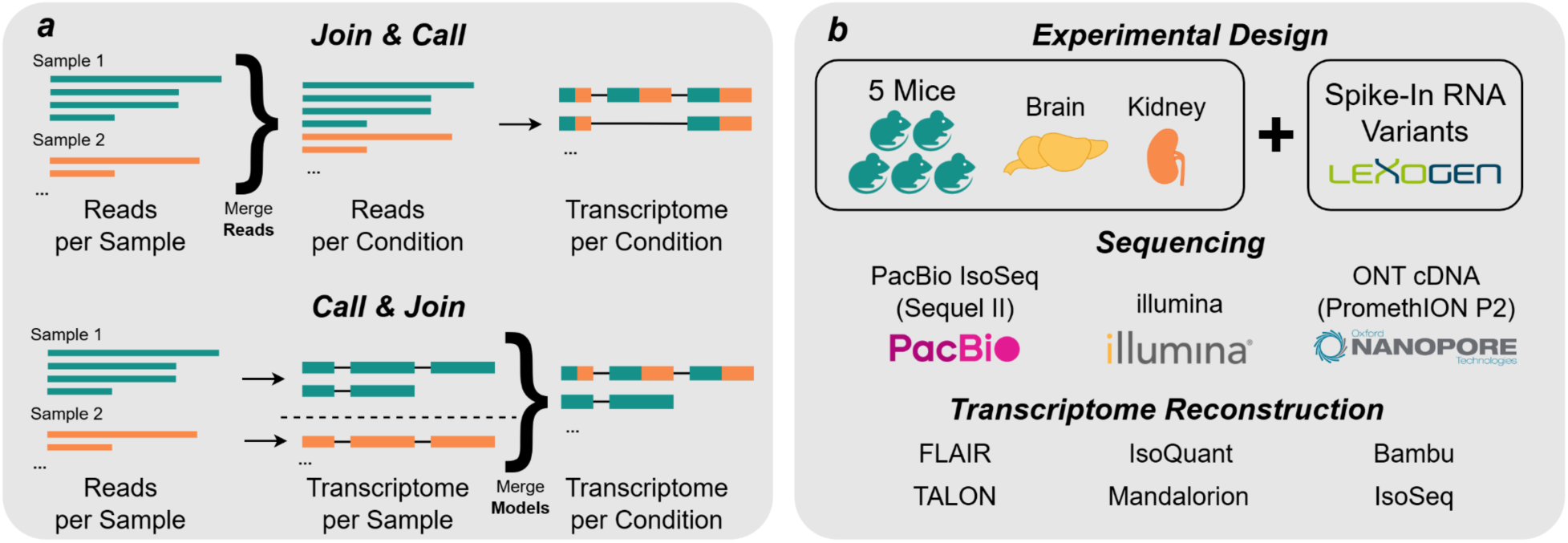
Overview of multi-sample combination strategies and experimental design. **a)** The Join and Call (J&C) strategy first merges reads from all replicates. Then, transcriptome reconstruction is performed, and eventually each sample is requantified using the joint transcriptome. Conversely, the Call and Join C&J strategy first performs transcriptome reconstruction for each replicate individually. Then, the resulting sets of transcript models are merged. **b)** Experimental set up to evaluate J&C and C&J strategies. Brain and kidney tissues from 5 mice were combined with Spike-In RNA Variants before sequencing with Pacific Biosciences (PacBio), illumina, and Oxford Nanopore Technologies (ONT). Six transcript reconstruction algorithms were evaluated: FLAIR, IsoQuant, Bambu, TALON, Mandalorion, and IsoSeq.

Here, we formally define and evaluate these two strategies by comparing J&C and C&J across six widely used transcript reconstruction tools applied to replicated lrRNA-seq datasets from mouse brain and kidney, generated with both PacBio and ONT technologies (**Figure 1b**). We show that strategy choice influences the discovery of rarely occurring novel transcripts, the detection of sample-specific transcripts, and the computational efficiency of the reconstruction. Furthermore, we investigate whether these effects are consistent across tissues and sequencing platforms, providing insight into the generalizability of strategy choice. Capturing such differences is particularly relevant in biological contexts where transcript usage is highly diverse, such as development, cellular differentiation or disease, and/or where rare isoforms can be functionally important^24–27^. By establishing these paradigms and quantifying their trade-offs, our study provides a conceptual and practical framework for multi-sample transcriptome reconstruction, guiding best practices in the era of increasingly large-scale lrRNA-seq studies.

## Results

To evaluate the J&C and C&J strategies for handling biological replicates in lrRNA-seq data, we selected six transcript reconstruction tools. Transcript model reconstruction is central to isoform discovery and quantification, yet different tools employ diverse algorithmic approaches and therefore exhibit distinct strengths and weaknesses. The LRGASP consortium highlighted that both sequencing platform and analysis tool strongly influence transcriptome definition, and that no method could reconstruct complete transcript models with high accuracy^11^. In their final recommendations, IsoQuant^5^, FLAIR^6^, and Bambu^7^ were favored for building sample-specific transcriptomes in well-annotated organisms with few novel isoforms, while Mandalorion^9^ and FLAIR were recommended for their sensitivity to detect lowly expressed or rare transcripts. Based on these guidelines, we selected IsoQuant, FLAIR, Bambu, and Mandalorion for evaluation, along with TALON^8^, the tool recommended by the ENCODE consortium^28^, and IsoSeq (https://isoseq.how/), the PacBio-supported pipeline. We further use TAMA Merge^10^ in order to combine transcript annotations in the C&J strategy.

Among these, IsoSeq uniquely performs reference-free transcript inference from PacBio HiFi reads, whereas the other five are reference-based methods. IsoQuant implements a graph-based approach, constructing splice graphs to represent the splicing landscape. FLAIR and Mandalorion are cluster-based methods that group reads by shared splice junctions or chains after genome alignment. Bambu and TALON apply class-based models, categorizing reads by splice-junction patterns to assign isoforms as known, novel, or ambiguous. Because IsoSeq often produces a large number of low-quality novel isoforms, we consistently applied SQANTI3-filter after IsoSeq reconstruction to improve transcript model quality, as recommended^29^.

To test whether J&C and C&J yield consistent outcomes across biological contexts and sequencing platforms, we employed a replicated lrRNA-seq datasets from two mouse tissues (brain and kidney) sequenced with both ONT (R9 flowcells) and PacBio (IsoSeq SMRTbell) platforms. Each tissue was collected from the same five mice, and complementary Illumina short-read sequencing was performed for all samples. In addition, synthetic spike-in RNA variant sequences (SIRVs) were included in all experiments to provide ground truth references^30^. Transcript reconstruction tools were run using their recommended parameters for ONT or PacBio data, with Illumina reads incorporated when suggested by the respective tool guidelines. For tools restricted to PacBio data (IsoSeq and Mandalorion), analyses were limited to PacBio datasets, while all other tools were evaluated on both platforms.

### Transcriptome Diversity and Isoform Counts

To evaluate how J&C and C&J influence transcriptome diversity, we first quantified the number of discovered transcripts by each of the 6 transcript reconstruction tools and then classified them into structural categories using SQANTI3 (**Figure 2**). According to SQANTI3 classification, full splice match (FSM) transcripts have the same splice junctions as annotated isoforms, while incomplete splice match (ISM) transcripts align to a known isoform but lack one or more splice junctions at one or both ends. Novel in catalog (NIC) transcripts use annotated splice sites in new combinations, whereas novel not in catalog (NNC) transcripts contain at least one unannotated splice site. Additional categories capture more divergent models: Genic Genomic transcripts overlap a known gene but extend beyond its annotated boundaries; Antisense transcripts align in the opposite orientation to annotated genes; Fusion transcripts span exons from two distinct genes; Intergenic transcripts map outside annotated genes; and Genic Intron transcripts originate entirely from within annotated introns.

**Figure 2:**
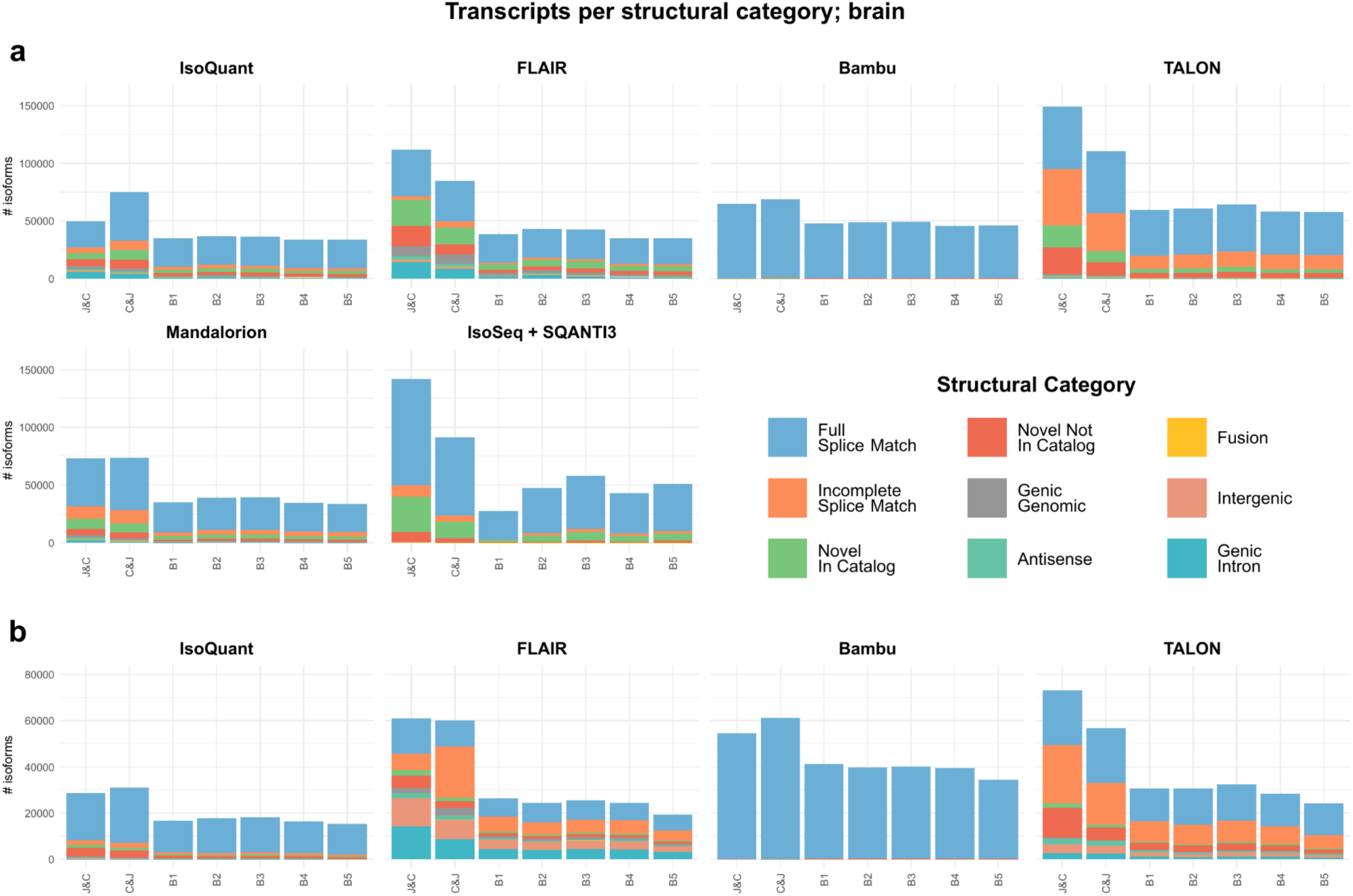
Transcripts per structural category in brain tissue. Transcriptome diversity as measured by SQANTI3 structural categories across combination strategies and individual samples and across transcriptome reconstruction tools in **a)** PacBio and **b)** ONT data. JC: Join & Call approach, C&J: Call and Join approach. B1-5. Mouse Brain samples 1 to 5.

In brain PacBio data, we observed a tool-dependent effect on the number of isoforms recovered between the two strategies (**Figure 2a**). For tools we consider more “permissive” in transcriptome reconstruction, such as FLAIR, TALON, and IsoSeq, the J&C strategy generally identified a greater number of isoforms, particularly novel ones. This outcome is anticipated, as pooling evidence from individual samples in J&C is expected to increase the confidence in calling lowly-expressed novel isoforms compared to analyzing samples independently in C&J. However, for IsoQuant on PacBio data, the opposite trend was observed, with C&J recovering more isoforms than J&C. We hypothesize that this behaviour may be related to IsoQuant’s splice site correction mechanism, which may cause reads to be assigned to distinctly structured transcript models in different samples and, therefore, combination strategies. Therefore, when samples are run in isolation, similar reads may be assigned to different isoforms, increasing the number of isoforms reported in C&J as compared to J&C, where the same reads may be pooled into a single isoform. Conversely, Bambu and Mandalorion appeared more robust to the choice of strategy when evaluated solely by the number of isoforms recovered. We hypothesize that this may be due to their reliance on reference annotations. When examining ONT data, the effects of combination strategy on isoform counts were less pronounced (**Figure 2b**). Further, ONT data generally resulted in a lower number of discovered isoforms. We hypothesize that this was, despite generally greater numbers of total reads, due to a notably higher fraction of unaligned reads, and shorter read lengths (**Supplementary Figure 1**). Differences between the structural categories of isoforms discovered in the two technologies were also reflected in the structural categories of reads according to analysis with SQANTI-Reads^31^. While the differences between strategies observed in IsoQuant, TALON, and Bambu remained similar, FLAIR showed little difference in the number of reported isoforms between the two strategies.

Next, we examined the SQANTI3 structural categories detected in each strategy. For FLAIR on ONT data, J&C recovered fewer ISM transcripts and more novel (NNC, NIC) transcripts compared to C&J, which showed a notable proportion of ISMs. This differed from the observations on PacBio data, where we did not observe an increase in ISMs in C&J. We hypothesize that this difference in the relative abundance of ISMs may be due to FLAIR’s definition of transcript ends benefitting from access to the increased amount of evidence provided by the J&C strategy, particularly on ONT data. For IsoQuant on ONT data, the previously discussed effect of differential splice junction calling across samples seemed less prominent, which might be attributable to IsoQuant’s wider splice junction correction window (6bp) for ONT data compared to PacBio (4bp). We found these observations to be consistent in kidney (**Supplementary Figures 2 and 3**).

**Figure 3:**
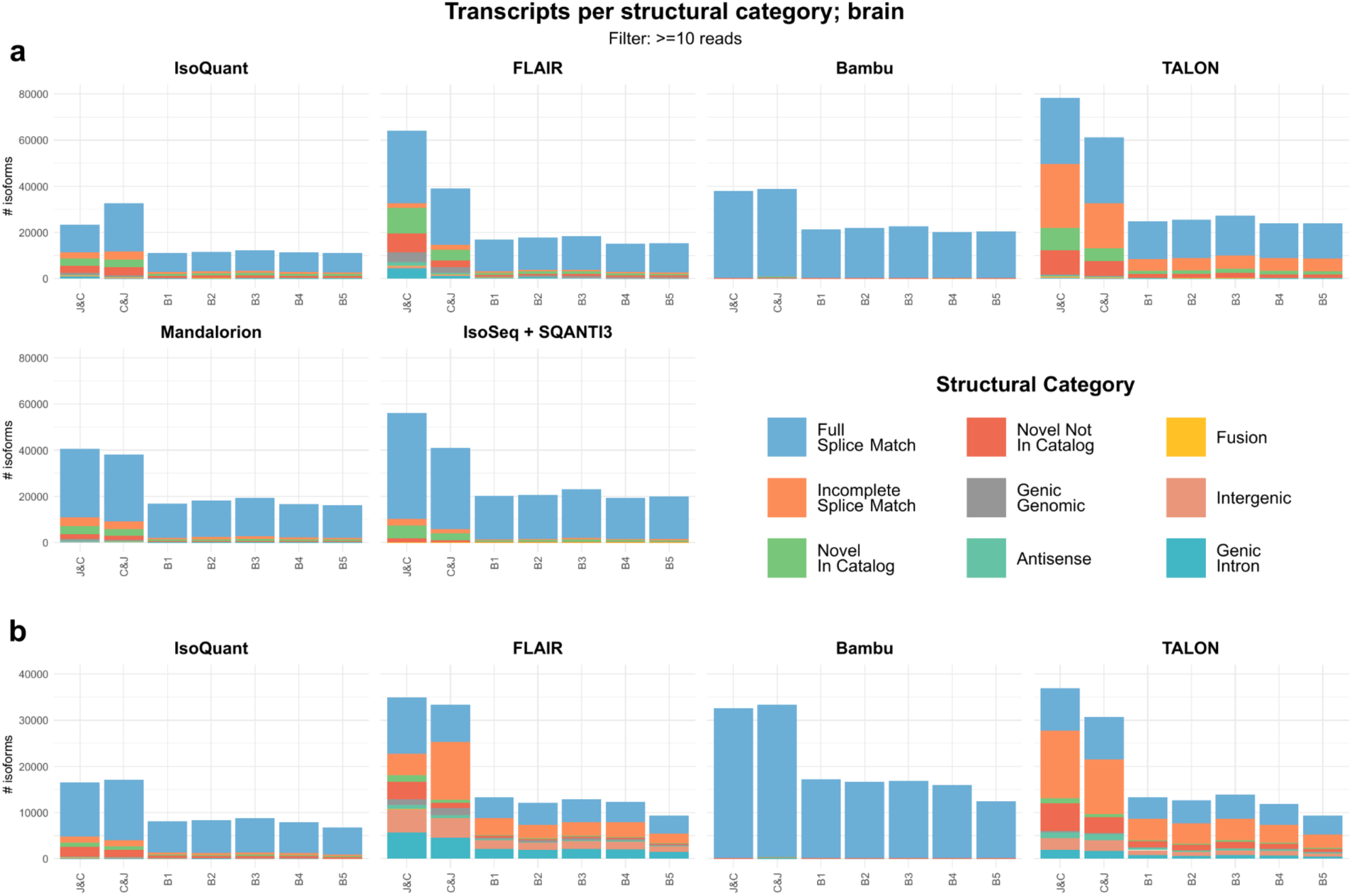
Transcripts with >= 10 reads per structural category in brain tissue. Transcriptome diversity as measured by SQANTI3 structural categories across combination strategies and individual samples and across transcriptome reconstruction tools in **a)** PacBio and **b)** ONT data. Filtered to transcripts with a minimum expression of 10. JC: Join & Call approach, C&J: Call and Join approach. B1-5. Mouse Brain samples 1 to 5.

Based on the analysis of transcriptome diversity and isoform counts, we conclude that the impact of J&C and C&J strategies on the number and structural categories of discovered transcripts is highly dependent on the transcript reconstruction tool and the sequencing platform used to generate the data.

### Impact of Expression Filtering on Transcriptome Diversity

After transcriptome reconstruction and quantification, analysis workflows typically curate the resulting transcriptome annotations by filtering out transcripts with low expression levels. To test how expression filtering might influence the identified transcriptome diversity, we applied expression filters of 5, 10 and 20 supporting reads and investigated the transcriptome composition across combination strategies, reconstruction tools and sequencing platforms, based on SQANTI3 structural categories (**Figure 3 and Supplementary Figures 4 and 5**).

Overall, the main patterns of transcriptome diversity and isoform counts remained robust after filtering, and the relative differences between strategies and tools were largely unchanged. The main effect of applying expression filters of 5, 10 and 20 supporting reads was a progressive reduction in the number of novel transcripts, while annotated transcripts were comparatively unaffected. This is a direct result of novel transcripts being generally more lowly expressed (**Supplementary Figure 9**). These observations were consistent in the kidney dataset (**Supplementary Figures 6, 7, 8, and 10**). Taken together, expression filtering within the ranges tested here, did not substantially alter the comparative behavior of combination strategies or reconstruction tools, but it reduced the discovery of lowly expressed and novel isoforms.

We further examined the number of reads supporting each transcript according to their structural category (**Supplementary Figure 9**), which revealed that the vast majority of reads were assigned to known (reference-annotated) transcripts, with novel transcripts generally being relatively lowly expressed. Interestingly, we observed strong differences in the number of unassigned reads across data types, with our ONT data exhibiting more unassigned reads (**Supplementary Figures 1 and 2**). Furthermore, for some tools (FLAIR, TALON, IsoSeq with PacBio data; FLAIR with ONT data), J&C was capable of assigning more reads than C&J, aligning with the premise that pooled evidence enhances the ability to call lowly-expressed novel transcripts with greater confidence, thereby enabling more reads to be assigned. These findings were largely consistent in kidney (**Supplementary Figure 10**).

### Intersection of Unique Junction Chains

To further understand how the J&C and C&J strategies affect not just the number of detected transcripts, but their identity, we compared their transcript models across samples and combination strategies. Since definition of transcript start sites (TSS) and transcript termination sites (TTS) is challenging in lrRNA-seq^29^, we compared Unique Junction Chains (UJCs; **Figure 4a for PacBio, Figure 4b for ONT**). We define UJCs as sets of splice junctions building the backbone of reported transcript models, while disregarding the TSS and TTS. Therefore, multiple transcripts which only differ in their start and termination sites (within the same terminal exon) will be represented by the same UJC, whereas transcripts with differences at their splice sites will be represented by different UJCs.

**Figure 4:**
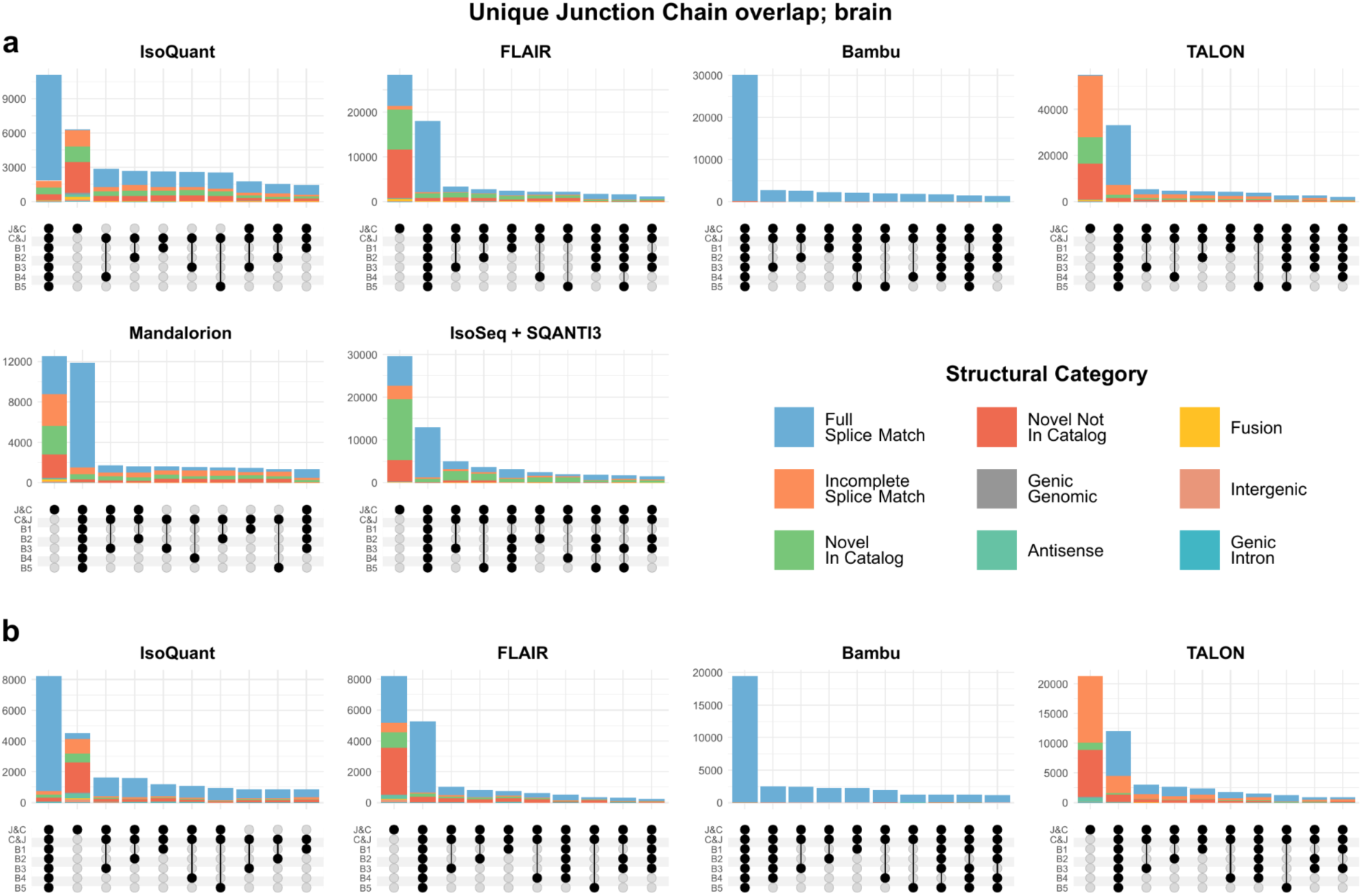
Unique Junction Chain intersections in brain tissue. 10 largest intersections of unique junction chains (UJCs) across combinations strategies and individual samples per transcriptome reconstruction tool for **a)** PacBio and **b)** ONT data. JC: Join & Call approach, C&J: Call and Join approach. B1-5. Mouse Brain samples 1 to 5.

For some tools (IsoQuant, Bambu), the largest intersection of UJCs were those found across all individual samples and both combination strategies, generally representing highly-expressed reference transcripts, indicating that these tools report fewer novel and/or sample-specific transcripts. Conversely, other tools (FLAIR, TALON, Mandalorion, IsoSeq) displayed the largest intersection as UJCs found only in the J&C strategy, followed by those found in all samples and strategies. This suggests that when evidence is combined in J&C, these tools are notably more proficient at discovering additional novel UJCs than in the C&J strategy. Furthermore, in the top 10 UJC intersections, some tools (FLAIR, Bambu, TALON, IsoSeq) showed no UJCs unique to the C&J strategy, meaning that nearly all UJCs discovered in C&J could also be found in J&C. This indicates a higher degree of stability or invariability to sample-specific changes in transcript definition for these tools. However, IsoQuant and Mandalorion did exhibit intersections of UJCs recovered in C&J but not in J&C. For IsoQuant, these C&J-exclusive intersections further support the notion that IsoQuant makes distinct transcript definition decisions across individual samples. For Mandalorion, we hypothesize that the C&J-exclusive UJCs observed in the upset plots can primarily be attributed to its default relative expression threshold of 1% of gene expression for reporting novel transcripts. These findings were consistent in kidney, except for Mandalorion, where the largest intersection switched from J&C-exclusive transcripts to UJCs expressed in all samples and strategies (**Supplementary Figure 11**). As the two intersections were similar sizes, this relatively small difference may be explained by the lower number of isoforms discovered in kidney (**Supplementary Figure 3**). Taken together, we conclude that for tools other than IsoQuant and Mandalorion, J&C results in a notable increase in the number of UJCs compared to C&J, and for all tools except for Bambu, J&C demonstrates a notable number of additional novel UJCs.

### Discovery of UJCs across samples

Building on the analysis of UJCs presented in Figure 4, we further investigated the discovery patterns of UJCs across varying numbers of samples.

Initially, we examined the number and structural categories of UJCs identified across 1, 2, 3, 4, or all 5 biological replicates (**Figure 5a for PacBio, Figure 5b for ONT**). Interestingly, across tools, the majority classes were UJCs present in only one sample or in all five samples, although the relative proportions varied depending on the transcriptome reconstruction method.

**Figure 5:**
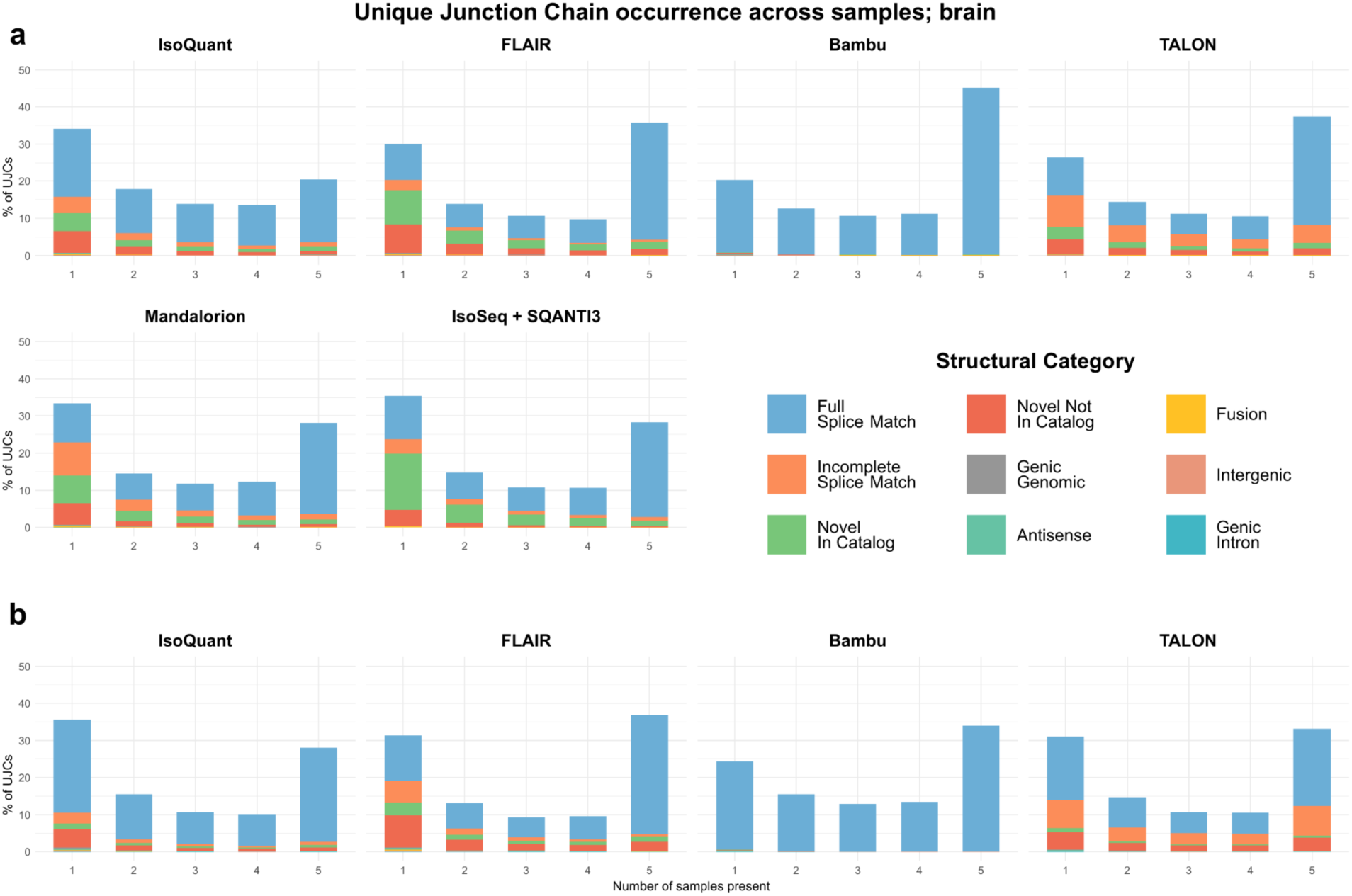
Distribution of unique junction chains (UJCs) across an increasing number of samples in brain tissue. Distribution of UJC occurrence across number of samples by structural category per transcriptome reconstruction tool for **a)** PacBio and **b)** ONT data of mouse brain tissue. The Y-axis represents the percentage of UJC observed in the indicated replicate count relative to the total identified UJC.

Subsequently, by considering the number of reads supporting UJCs (**Figure 6a for PacBio, Figure 6b for ONT**), we confirmed that for most tools, the vast majority of reads were assigned to UJCs detectable in all five samples. This outcome is expected, given that the majority of reads correspond to highly-expressed reference transcripts (**Supplementary Figures 9 and 10**). A notable exception to this pattern is the performance of IsoQuant, particularly with PacBio data, where a discernibly larger proportion of reads were assigned to UJCs identified in only a subset of samples. As reported in the section “Transcriptome diversity and isoform counts”, this pattern likely suggests that IsoQuant, when performing isoform identification on individual samples as part of the C&J strategy, may generate slightly different splice junction definitions compared to when the same reads are combined in the J&C strategy. These findings were consistent in kidney (**Supplementary Figures 12 and 13**).

**Figure 6:**
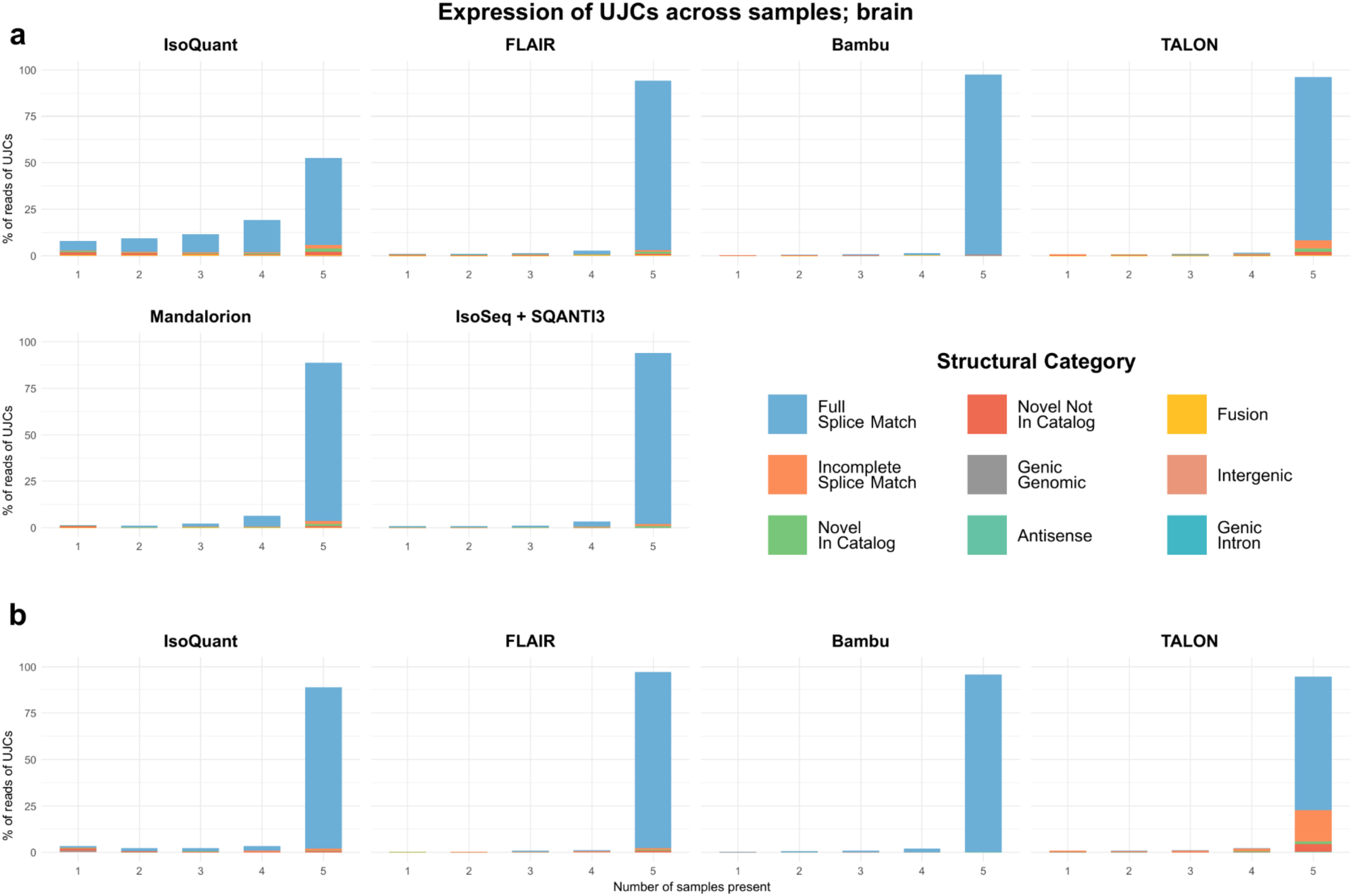
Distribution of expression of unique junction chains (UJCs) across an increasing number of samples in brain tissue. Distribution of expression of UJC occurrence across number of samples by structural category per transcriptome reconstruction tool for **a)** PacBio and **b)** ONT data of mouse brain tissue. The Y-axis represents the percentage of UJC observed in the indicated replicate count relative to the total identified UJC.

### Evaluation with Spike-in Controls and TUSCO

To comprehensively evaluate the performance of the combination strategies and the various isoform identification tools, we leveraged external spike-in RNA variants^30^ (SIRVs) and TUSCO^32^ (**Figure 7 for SIRVs, Figure 8 for TUSCO**). SIRVs are defined synthetic RNA molecules which are added to extracted RNA samples before library preparation, to serve as a control of technical errors introduced in library preparation and sequencing. TUSCO is a curated reference set of genes lacking alternative isoforms used as an endogenous control of transcriptome reconstruction workflows. The metrics utilized for this assessment, which are applied to both SIRVs and TUSCO, include sensitivity, positive detection rate, non-redundant precision, redundant precision, redundancy, and false discovery rate (see Methods).

**Figure 7:**
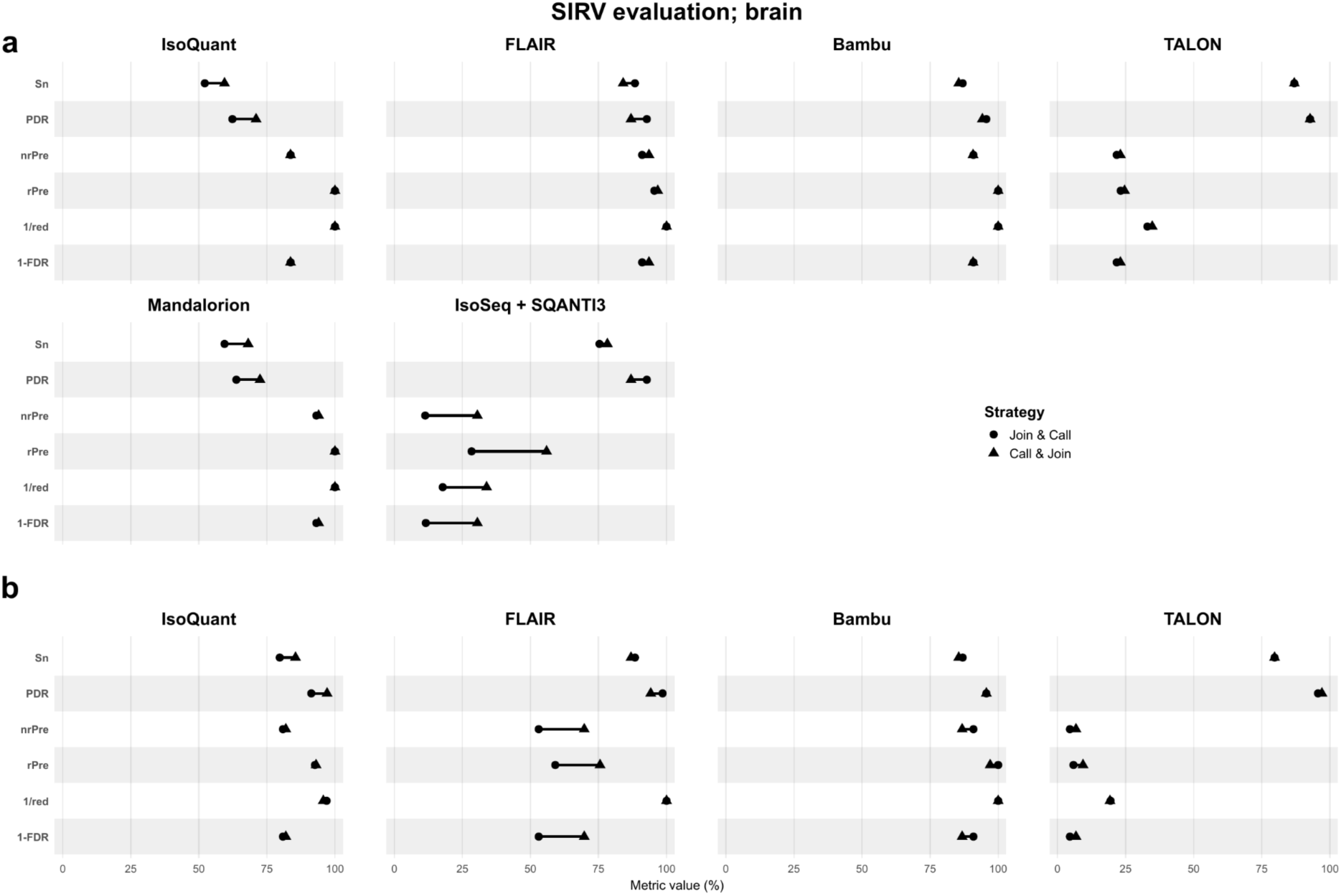
SIRV evaluation in brain tissue. Sensitivity (Sn), positive detection rate (PDR), non-redundant precision (nrPre), redundant precision (rPre), redundancy (red) and false discovery rate (FDR) for SIRV analysis comparing the combination strategies per transcriptome reconstruction tool for **a)** PacBio and **b)** ONT data of mouse brain tissue. In order for all metrics to fit a 0-100% scale where higher is better, redundancy and false discovery rate have been converted to 1/red and 1-FDR.

**Figure 8:**
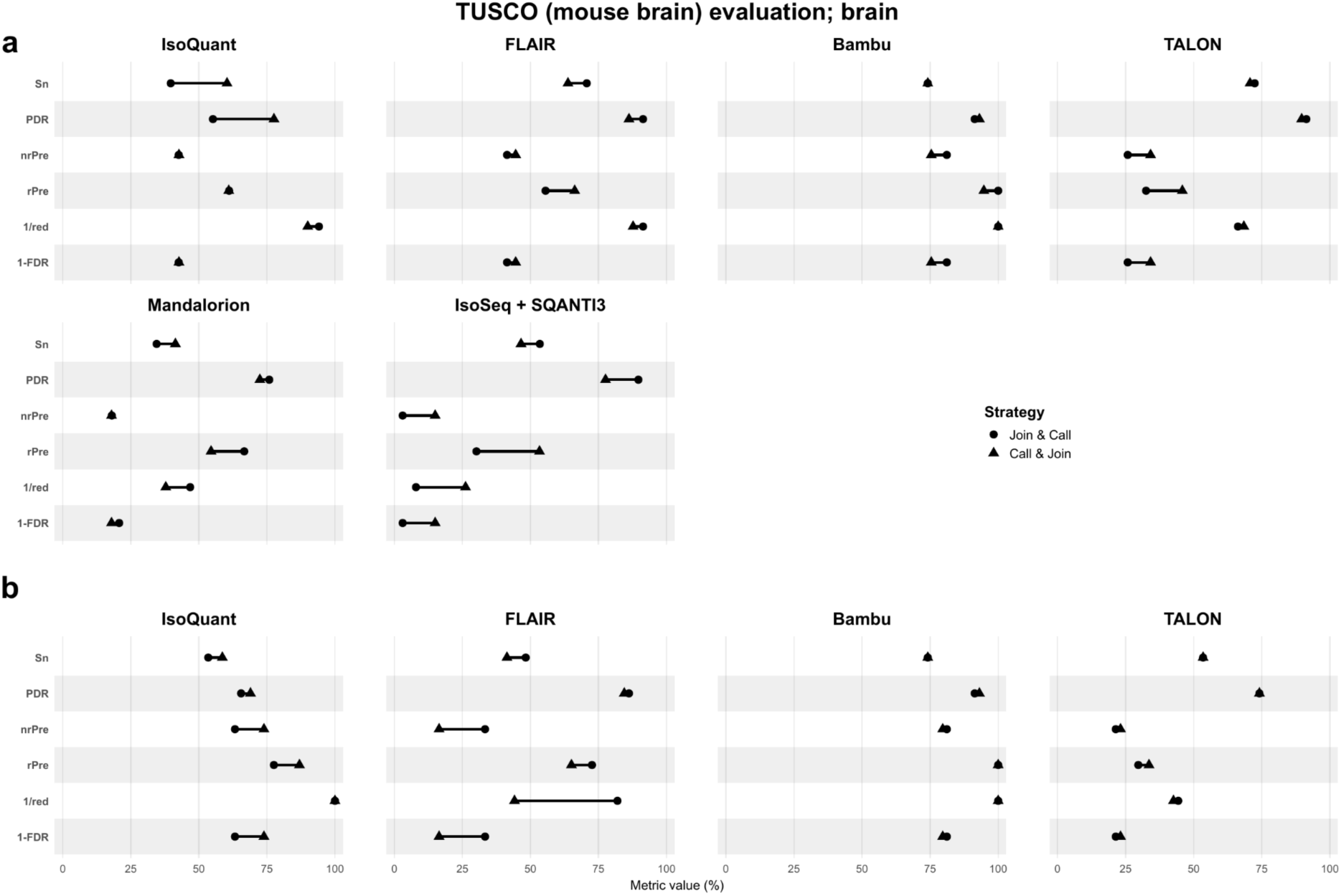
TUSCO (mouse brain) evaluation in brain tissue. Sensitivity (Sn), positive detection rate (PDR), non-redundant precision (nrPre), redundant precision (rPre), redundancy (red) and false discovery rate (FDR) for TUSCO (mouse brain) analysis comparing the combination strategies per transcriptome reconstruction tool for **a)** PacBio and **b)** ONT data of mouse brain tissue. In order for all metrics to fit a 0-100% scale where higher is better, redundancy and false discovery rate have been converted to 1/red and 1-FDR.

Our analysis revealed that the evaluation of outcomes based on TUSCO and SIRVs did not always align. Generally, SIRVs analyses resulted in better performance metrics, and greater similarity between C&J and J&C than TUSCO, except for precision-related metrics in TALON, where SIRVs indicated poorer performance. We hypothesize these differences reflect the generally lower expression values and higher length of TUSCO transcripts with respect to SIRVs, with TUSCO better recapitulating transcriptome characteristics (**Supplementary Figures 18 and 19**). Because TUSCO is an internal standard derived from the samples, it captures internal variation, expression level dynamics and transcript lengths distributions, providing a more biologically representative benchmark. These results suggest that evaluations based solely on SIRVs may overestimate the performance of transcriptome reconstruction tools^32^.

Beyond these differences, performance comparisons between C&J and J&C were broadly consistent when employing SIRVs or TUSCO. In agreement with observations from the analysis of transcripts, the superior performance of either J&C or C&J strategies on a given metric was dependent on the specific tool and dataset being evaluated. Both Bambu and TALON showed very similar performance for the C&J and J&C approaches in both PacBio and ONT datasets, with only a slightly higher non-redundant precision observed for TALON with the C&J strategy in PacBio data. IsoQuant performed equally well or better with C&J than with J&C across all performance metrics and for both sequencing technologies. This may be attributed to its previously discussed tendency to collapse isoforms more extensively under the J&C strategy. In contrast, for FLAIR, precision-related metrics were higher on the ONT dataset when applying the J&C strategy. We reason that this is due to FLAIR’s constraint of reporting a single TSS and TTS per unique junction chain in J&C, whereas C&J different TSS/TTS might be called for each sample that remained separated after TAMA that uses a <= 50nt offset window for collapsing. This negatively affects transcript redundancy and hence precision. Finally, the IsoSeq+SQANTI3 pipeline showed better sensitivity and PDR with J&C but higher precision-related metrics with C&J. These results were largely consistent in kidney samples (**Supplementary Figures 15 and 16**), as well as when using TUSCO’s tissue-nonspecific isoform sets (**Supplementary Figures 14 and 17**).

Taken together, these results further indicate that the relative performance of the C&J and J&C strategies varied mainly with the reconstruction tool and sequencing platform, rather than showing a consistent trend across datasets. IsoQuant generally favored C&J across metrics, whereas FLAIR and IsoSeq+SQANTI3 showed metric-specific shifts between strategies, particularly between PacBio and ONT datasets. While some discrepancies were observed between SIRV- and TUSCO-based evaluations, the overall patterns of tool- and technology-dependent behavior remained largely comparable.

### Computational Resource Requirements

In order to assess the computational resource requirements of the two combination strategies, primarily execution time and memory usage, we recorded elapsed time, CPU time, and maximum memory usage of each process (see Supplementary Table B).

As expected, both CPU time and memory usage varied widely by transcriptome reconstruction tool, although the sample-combination strategy also showed notable impact (**Figure 9a for PacBio, Figure 9b for ONT; Supplementary Figure 20 for kidney**). FLAIR, TALON, and Mandalorion had the highest runtimes in both J&C and C&J, while TALON and Mandalorion had the highest memory usage, particularly with J&C. Lowest runtimes overall, regardless of combination strategy, were achieved by Bambu and IsoQuant, while lowest memory consumption was achieved by IsoQuant and FLAIR.

**Figure 9:**
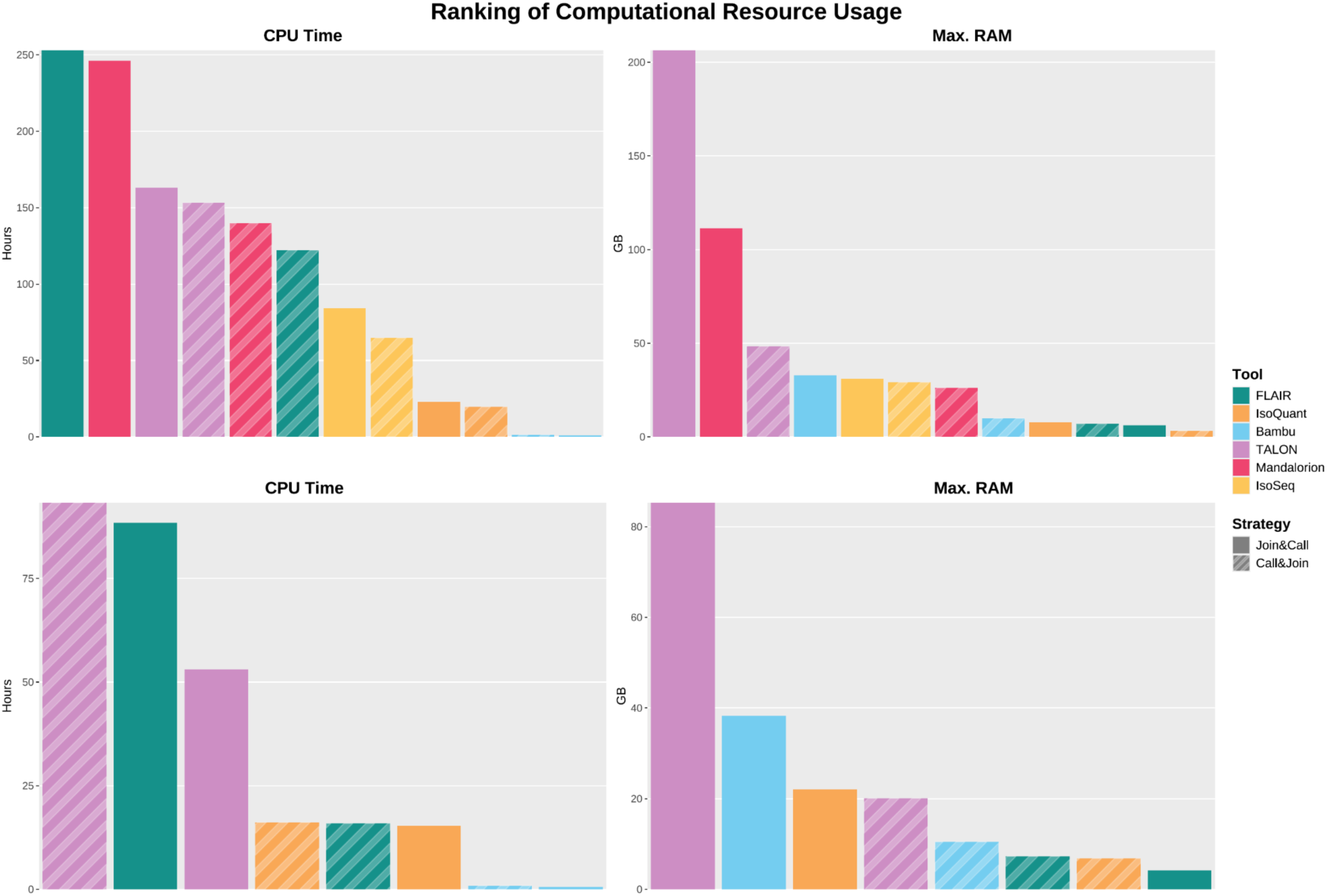
Ranking of CPU Time and Maximum RAM Usage in brain tissue. Ranking of transcriptome reconstruction tools and combination strategies by their required CPU Time in hours and maximum memory usage in gigabytes in **a)** PacBio and **b)** ONT data of mouse brain tissue. For C&J, the CPU time measurements are the sum of the individual processes for each sample plus the process for TAMA Merge, while the maximum memory is the maximum of the individual processes. For J&C, wherever processes were split (e.g. by chromosome or for identification and quantification), the measurements were aggregated in the same way.

For Mandalorion, FLAIR, IsoSeq, and IsoQuant, both CPU time and memory usage were lower under C&J than J&C. This was expected as the same data is split into a larger number of processes. For TALON (except for PacBio data of brain tissue) and Bambu, CPU time was lower in J&C than in C&J. We hypothesize that this may point to a larger proportion of time spent on overhead tasks. With the exception of FLAIR, memory usage of all tools was lower in C&J than in J&C. FLAIR with J&C had a lower than expected memory usage due to splitting the *FLAIR collapse* process by chromosome, as recommended for large datasets (see Methods).

## Discussion

In this study, we address a fundamental and timely question in long-read transcriptomics: *how should lrRNA-seq multi-sample datasets be combined to reconstruct a biologically meaningful transcriptome?* This question is becoming increasingly important as long-read sequencing shifts from low-throughput, single-sample applications toward large studies with many biological replicates. Previously, pooling reads across samples has been a common approach for defining consensus transcriptomes, especially when throughput was limited. However, as higher read yields make multi-sample designs standard, pooling is not always practical or scalable, and separate per-sample analyses followed by transcriptome merging is an appealing alternative. However, these alternatives have not been systematically assessed. Here, by formally defining and benchmarking the *Join & Call* (J&C) and *Call & Join* (C&J) paradigms across two tissues, two sequencing platforms, and six reconstruction tools, we demonstrate that the choice of strategy exerts a measurable and tool-dependent impact on transcript discovery, isoform structure, read assignment, and computational efficiency. These findings establish a conceptual framework for multi-sample long-read transcriptomics and provide practical guidelines for designing future large-scale studies.

A central conclusion of our work is that no single combination strategy universally outperforms the other. Instead, the relative strengths of J&C and C&J are strongly modulated by both algorithmic design and data characteristics. Tools such as FLAIR, TALON, and IsoSeq benefited substantially from evidence pooling in J&C, recovering a larger diversity of isoforms—particularly novel UJCs—than when samples were processed independently. By pooling reads across samples, J&C can call transcript models that would otherwise fall below per-sample detection limits, and can increase the number of reads successfully assigned to transcripts. This confirms the intuitive expectation that increased read depth across replicates enhances sensitivity to rare or lowly expressed isoforms. In contrast, IsoQuant consistently showed greater transcript diversity in C&J, largely due to its splice-site correction and graph-based inference, which can generate slightly distinct transcript models across samples and thus inflate diversity when processed independently. These opposite behaviors illustrate that transcript model definition is not solely determined by read data but also by tool-intrinsic heuristics regarding splice junction correction, clustering, and annotation reliance.

A key insight from our analysis is that relative expression thresholds used by tools such as IsoQuant and Mandalorion influence isoform discovery differently under the J&C and C&J strategies. These tools require not only that a transcript surpass an absolute read-count cutoff, but also that it represents a minimum fraction of its gene’s total expression (e.g., ≥2%). Under C&J, a lowly expressed novel transcript may exceed this relative threshold within individual samples and therefore be retained. In contrast, when reads are pooled in J&C, the gene’s total expression increases, causing the same transcript to fall below the relative threshold and go undetected (see Supplementary Table A). This mechanism accounts for a substantial portion of the strategy-specific differences we observed and illustrates how filtering criteria interact with multi-sample data structure in ways that are not always intuitive Additionally, although expression filtering is widely applied to mitigate model inflation, we find that filtering thresholds (5–20 supporting reads) do not materially alter the relative behavior of tools or combination strategies. Instead, filtering predominantly reduces novel transcript calls—more lowly expressed by nature—while leaving annotated isoforms largely intact. Thus, filtering can temper, but not eliminate, the tendency of J&C to recover more novel isoforms or the tendency of C&J (for certain tools) to generate more structurally heterogeneous transcript sets. This suggests that the choice of combination strategy should be made *prior* to filtering decisions, as filtering does not compensate for upstream methodological differences.

Using both SIRVs and TUSCO, we assessed performance in contexts with known ground truth. Although performance estimates differed between the two benchmarks—reflecting differences in transcript length, expression range, and biological complexity—the overall patterns of tool-specific sensitivity and precision trade-offs remained stable. Across metrics, strategy performance was rarely consistent across tools: J&C improved sensitivity for IsoSeq+SQANTI3 and, under ONT data, improved precision for FLAIR, while IsoQuant consistently achieved higher precision and sensitivity under C&J. This variability indicates that combination strategy influences not only the number of transcripts called but also their correctness, and that strategy selection should consider both tool behavior and the biological objectives of the study.

From a practical standpoint, computational requirements represent an important and often overlooked dimension of multi-sample long-read transcriptomics. As expected, C&J was markedly more computationally efficient, reducing both runtime and peak memory for most tools, because it distributes inference across samples and avoids constructing large global clustering structures. By contrast, J&C frequently required >100 GB of memory for tools such as TALON or Mandalorion. As lrRNA-seq studies continue to scale into dozens or hundreds of replicates, the feasibility of J&C may become limiting, particularly in resource-constrained environments. Thus, when discovery of rare isoforms is not the primary goal, C&J offers a practical alternative with substantially reduced computational cost.

Together, these results support a use-case–specific decision model for multi-sample lrRNA-seq. For studies prioritizing rare-isoform discovery, e.g. cell differentiation, disease-associated splicing, or fusion detection, J&C is advantageous, especially with inherently permissive tools such as FLAIR or IsoSeq+SQANTI3. However, we also strongly recommend that these approaches are followed by rigorous quality control, including –when possible– incorporating support by orthogonal data, to mitigate false discoveries. Conversely, for studies emphasizing quantification, differential transcript usage, or large cohort scalability, C&J with more conservative tools such as Mandalorion or Bambu provides more stable performance and dramatically lower computational burden. Tool choice interacts strongly with strategy choice, meaning that recommendations cannot be generalized across the field without considering the analytic pipeline in full.

Finally, the conceptual distinction between J&C and C&J highlights a broader point: multi-sample long-read transcriptomics is not yet standardized, and methodological decisions can have downstream consequences for isoform-level analyses, variant interpretation, and reference-free transcriptomics. We therefore advocate that future benchmarking efforts explicitly incorporate multi-sample scenarios and that sequencing consortia adopt standard reporting of combination strategy, filtering, and merge thresholds. As rapid improvements in read accuracy and throughput continue to expand the scale of lrRNA-seq projects, our framework provides a foundation for robust, reproducible, and biologically meaningful transcriptome reconstruction

## Methods

Five male C57BL/6J wild type mice were obtained from Jackson Laboratories and group housed with standard chow and tap water ad libitum at the University of Valencia animal facility (Servei Central de Suport a la Investigació Experimental - SCSIE, Burjassot). At 3 months of age, mice were anesthetized with 60 mg/kg pentobarbital (i.p.) and perfused intracardially with DEPC treated 0.9% saline to minimize blood contamination. Brains and kidneys were immediately collected and maintained in ice cold DEPC treated phosphate buffered saline at all times. The interval between tissue collection and RNA extraction was kept as short as possible and consistent across all samples and mice. Before RNA isolation, cerebellum, medulla and olfactory bulbs were removed from the brain samples. For kidney samples, suprarenal glands, ureters and the surrounding connective capsule were dissected away. All animal handling and experimental procedures were conducted in accordance with European Union 86/609/EEC and Spanish RD1201/2005 guidelines, following protocols approved by the Ethics Committee on Experimental Research of the University of Valencia.

### RNA isolation

Total RNA was extracted using the Maxwell® 16 LEV simplyRNA Purification Kit (Promega). Each brain hemisphere and kidney were homogenized separately in the kit’s homogenization buffer using a FastPrep system (MPBio) with 1.4 mm zirconium beads, 2 × 30 s pulses. Homogenates from the same mouse and tissue were combined before automated lysis and RNA extraction on a Maxwell 16 device (Promega), following the manufacturer’s instructions. Two independent extractions were performed per lysate that were finally pooled, yielding RNA representing ∼27% of the original tissue homogenate. RNA quantity and quality were assessed using the Qubit RNA BR Assay (ThermoFisher Scientific) and the Agilent TapeStation System (Agilent Technologies). Extracted RNA was spiked with Lexogen SIRV-Set 130 (∼3% of total RNA molecules sequenced), with brain samples receiving SIRV subset E0 and kidney samples SIRV subset E1.

### PacBio library preparation, sequencing and data preprocessing

cDNA synthesis and sample barcoding were performed using the NEBNext Single Cell/Low Input cDNA Synthesis and Amplification Kit, following the standard protocol. Barcoded cDNA samples were mixed into two pools: the first containing 3 brains and 3 kidneys from the same 3 mice, and the second containing 2 brains and 2 kidneys from the same 2 mice. Four libraries were generated for each pool using IsoSeq SMRTbell prep kit 3.0. A total of 16 SMRT Cells per pool were sequenced over 8 runs on a PacBio Sequel IIe at the University of Valencia (Servei Central de Suport a la Investigació Experimental – SCSIE, Burjassot).

Lima (v2.9.0) was used for demultiplexing and primer removal from HiFi unaligned BAM files. PolyA tails trimming and removal of concatemers was performed using isoseq refine. After converting BAM to FASTQ files using BamTools^35^, reads were mapped to the reference mm39 NCBI RefSeq genome^36^ using minimap2^37^ (v2.17) with options -ax splice:hq -uf.

### Nanopore library preparation, sequencing and data preprocessing

cDNA synthesis and sample barcoding were performed using the Nanopore SQK-PCB111-24 PCR-cDNA Barcoding Kit, following the standard protocol and generating 8 libraries per sample. The 8 libraries were then mixed and split into 8 aliquots. Eight rounds of sequencing were conducted on R9 flow cells (FLO-PRO002) using a Nanopore PromethION P2 at LongSeq applications.

Raw Pod5 files were demultiplexed using Dorado (v7.2.13) with the High-accuracy model, 450 bps, on the MinKNOW cloud. fastqc (v0.12.0) was used to verify read quality with a phred score above 20. FASTQ files per sample were merged, and stranded using Pychopper (v2.5.0). Full-length transcripts were mapped to the reference genome mm39 NCBI RefSeq, using minimap2 (v2.17) with options -ax splice -uf.

### Illumina library preparation, sequencing and data preprocessing

Directional cDNA synthesis and sample barcoding were performed using Illumina TruSeq Stranded mRNA Kit, following the standard protocol. After pooling the libraries, the samples were sequenced at 150x2 bp on an Illumina NovaSeq at Novogene.

Raw BCL files were automatically demultiplexed using bcl2fastq (v2.20) on the BaseSpace Sequence Hub. Quality control of FASTQ files was performed using fastqc (v0.12.0), and mapping to the reference genome mm39 NCBI RefSeq was performed using STAR^38^ (2.7.10b).

### Read-level analysis

Read-level analysis was performed with SQANTI-Reads (v5.3.6) to identify the structural categories of reads.

### Isoform Identification

We examined multiple different tools commonly used in the community for the identification of isoforms: the IsoSeq pipeline^39^, Mandalorion^9^, FLAIR^6^, TALON^8^, IsoQuant^5^, and Bambu^7^. All of these tools were used on the PacBio data, whereas all except IsoSeq and Mandalorion were also used on the ONT data, as IsoSeq is specifically designed for PacBio data and Mandalorion is designed for circular consensus sequence reads (PacBio or ONT with R2C2^40^). Unless otherwise stated, the tools were run with their default parameters.

### Join & Call

In J&C the reads of all samples were combined into a single file per tissue. The isoform identification tools were then run, unaware of which read originated from which sample. The end result of each tool was a joint transcriptome per tissue. Most examined tools allowed the use of such a transcriptome with the FASTQ files of the individual samples to determine transcript abundances for each sample.

### Call & Join

For C&J the isoform identification tools were run on each sample individually, completely independent of each other. The resulting sample-specific transcriptomes were then merged into a joint transcriptome per tissue using TAMA Merge^10^.

To configure TAMA Merge (commit 2fa3c30), we used thresholds of 50 base pairs for the transcription start and end sites (options *-a 50* and *-z 50*) and a strict threshold of 0 base pairs for splice junctions (*-m 0*). The rationale for the choice of these parameters was that the UJC of a transcript is central to transcript identity, while variation at the start and termination site could occur across different samples due to random variation. We also supplied the option *-d merge_dup* to merge duplicate transcript models. Furthermore, in the input file list (*-f*), we set the *cap_flag* as *capped* for all transcriptomes in order to avoid collapsing transcript ends beyond the set thresholds of 50 base pairs and used equal merge priorities of *1,1,1*.

### IsoQuant

We ran IsoQuant (3.6.3) with the *--fl_data* option for PacBio, and without it for ONT data.

### FLAIR

We ran FLAIR (v2.1.1) by first converting the alignment BAM format files to BED using FLAIR’s *bam2Bed12* program.We then ran *flair correct* utilizing both the reference transcriptome, as well as short-read junctions in the form of the *SJ.out.tab* output file of STAR, as recommended. Within J&C, we supplied these short-read junctions from all samples, whereas for C&J we only supplied short-read junctions obtained from the same sample.

We further ran *flair collapse* using the parameters *--stringent*, *--check_splice*, and *--annotation_reliant generate* as recommended by the documentation. In J&C, this step was performed by splitting the input file by chromosome and running each chromosome individually, as recommended for large datasets by the documentation.

Finally, we ran *flair quantify*. Here, for C&J, we used the *--stringent* and *--check_splice* parameters. However, they could not be run on J&C.

### Bambu

We ran Bambu (3.8.3) with default settings.

### TALON

To run TALON (6.0.1), we first initialized a database with *talon_initialize_database*. We further ran, with default parameters and as recommended by the documentation, the programs *transcriptclean* (2.1), *talon_label_reads*, *talon*, *talon_filter_transcripts*, *talon_create_GTF*, and finally *talon_abundance*.

### Mandalorion

We ran Mandalorion (v4.5.0) with the *-W ‘[SIRV,“basic”]’* parameter as recommended by the documentation to whitelist polyA-sites of SIRV and ‘basic’ transcripts.

### IsoSeq + SQANTI3

To run IsoSeq (v4.0.0), we first ran *isoseq cluster2*, followed by alignment using *pbmm2 align* with the parameters *--preset ISOSEQ --sort*. We then ran *isoseq collapse* with the parameter *--do-not-collapse-extra-5exons*.

In order to filter this highly-redundant initial transcriptome, we used SQANTI3 (v5.3.6) Quality Control and Filter, as recommended^29^. We provided FL-counts, short-read alignments and junctions, CAGE-peaks, polyA-peaks, and polyA-motifs to SQANTI3 QC. We further ran the SQANTI3 machine learning filter, for which we manually defined true positive and true negative sets for the training step. As true positives, we selected multi-exon FSM transcripts with only canonical splice junctions that also had support by both polyA- and CAGE-peaks. As true negatives, we selected multi-exon non-FSM transcripts with either at least one non-canonical splice junction, no CAGE-peak support, or no polyA-peak support. In order to avoid overfitting of the machine learning model, we then excluded the columns *all_canonical*, *within_CAGE_peak*, *within_polyA_site*, *dist_to_CAGE_peak*, and *dist_to_polyA_site* from the data.

### Post-processing

The above-described pipeline resulted in the creation of 7 transcriptomes per tissue (1 from J&C + 5 individual samples and 1 combined from C&J), per isoform identification tool. As we found some tools to report transcripts despite an expression of 0, we further applied a filter to limit our analyses to transcripts with an expression of at least 1 read.

To evaluate our results, we ran SQANTI3^29^ (5.3.5) Quality Control on each resulting transcriptome to determine the structural categories of the identified isoforms in comparison to the mm39 NCBI RefSeq transcript annotations, modified to include the SIRVs. We used TUSCO version 0.99. Metrics used for the evaluation of SIRVs and TUSCO isoforms were calculated as described in TUSCO^32^. For clarity and self-containment, we briefly summarize the calculation of these metrics here:

True positives (*TP*) are defined as transcripts matching the intron-exon structure of a reference TUSCO gene, as well as having their Transcription Start Site (TSS) and Transcription Termination Site (TTS) within +/-50bp (SQANTI3 structural category full-splice_match (FSM), subcategory reference_match).

Partial true positives (*PTP*) are defined as FSM or incomplete-splice_match (ISM) transcripts with deviations at their TSS and/or TTS.

False negatives (*FN*) are defined as TUSCO genes for which no transcripts are observed. Further, let *G* be the number of TUSCO genes, *TP_g_* the number of TUSCO genes with at least one *TP*, *D* the number of TUSCO genes with at least one *TP* or *PTP* transcript, and *T_f_sm + ism* the number of FSM plus ISM transcripts.

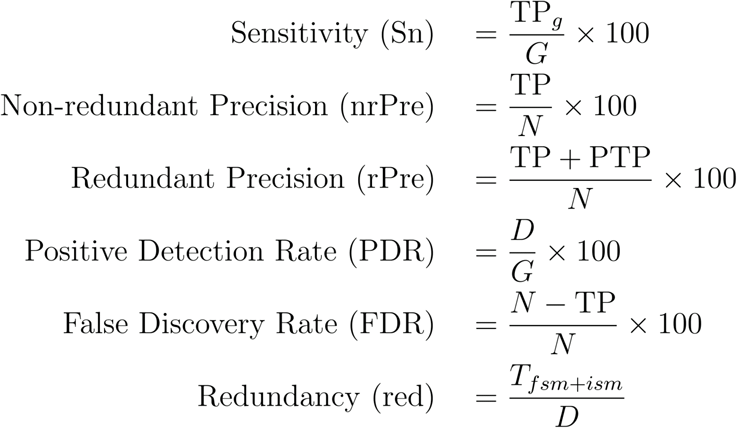

Finally, we investigated the structural categories of UJCs.

### Computational resource usage

Transcript identification and post-processing analyses were performed using a custom nextflow pipeline, with individual jobs running on a SLURM cluster. Using the SLURM system, elapsed time, CPU time, and maximum memory usage of each individual job were recorded (see Supplementary Table B). In order to compare usage of computational resources of different tools and combination strategies, data was aggregated in the following manner:

For C&J, each sample was run in an individual job, followed by TAMA Merge. The resource usage was calculated by taking the sum of elapsed times and, likewise, CPU times of all samples and the TAMA Merge job, and taking the maximum of the memory usage.

For J&C, where jobs were separated e.g. in identification and quantification or, for FLAIR, by chromosome, the metrics were aggregated in the same manner.

## Data availability

All data generated in this study have been submitted to the European Nucleotide Archive (ENA; https://www.ebi.ac.uk/ena/browser/home). Mice brain and kidney data generated using PacBio, ONT and Illumina sequencing are accessible under accession numbers PRJEB85167 and PRJEB94912.

## Code availability

To facilitate future evaluation of J&C vs C&J strategies with additional datasets, transcript reconstruction tools and their configurations, we provide a nextflow^34^ pipeline that enables systematic evaluation of the multi-sample combination strategies. It is available at https://github.com/ConesaLab/join_and_call_paper

## Acknowledgements

This project has received funding from the European Union’s programme Horizon Europe under the Marie Skłodowska-Curie grant agreement Number 101072892 and 101149931, the Spanish MICINN (PID2020-119537RB-I00 and FPU21/01597), MICIU/AEI (PID2023-152976NB-I00) and the National Institutes of Health (1R21HG011280-01).

The computations were performed on the high performance computing cluster Garnatxa at the Institute for Integrative Systems Biology (I^2^SysBio), I^2^SysBio is a mixed research centre formed by University of Valencia (UV) and Spanish National Research Council (CSIC).

## Author contributions

F.J. performed analysis on PacBio and ONT data with FLAIR, TALON, IsoQuant, Bambu, and SQANTI3. A.P. performed analysis on PacBio data with IsoSeq, SQANTI3 Filter, Mandalorion, and Bambu. J.M-T. performed initial analysis of PacBio data with FLAIR. S.C. performed initial analysis on PacBio data with Mandalorion and Bambu. F.J., A.P. and C.M. performed data pre-processing. F.J., A.P., C.M. and A.C. analysed results. J-M.M-R, I.F. and L.F. generated the datasets. F.J., C.M. and A.C. wrote the manuscript. S.G. supervised F.J.; A.C., C.M. and A.P. supervised the project.

## Ethics declarations

A.C. has received in-kind funding from Pacific Biosciences for library preparation and sequencing. A.C. and F.J. collaborate with Oxford Nanopore in the Marie Skłodowska-Curie Actions Doctoral Network project LongTREC.

## Supplementary Information

**Supplementary Figure 1:**
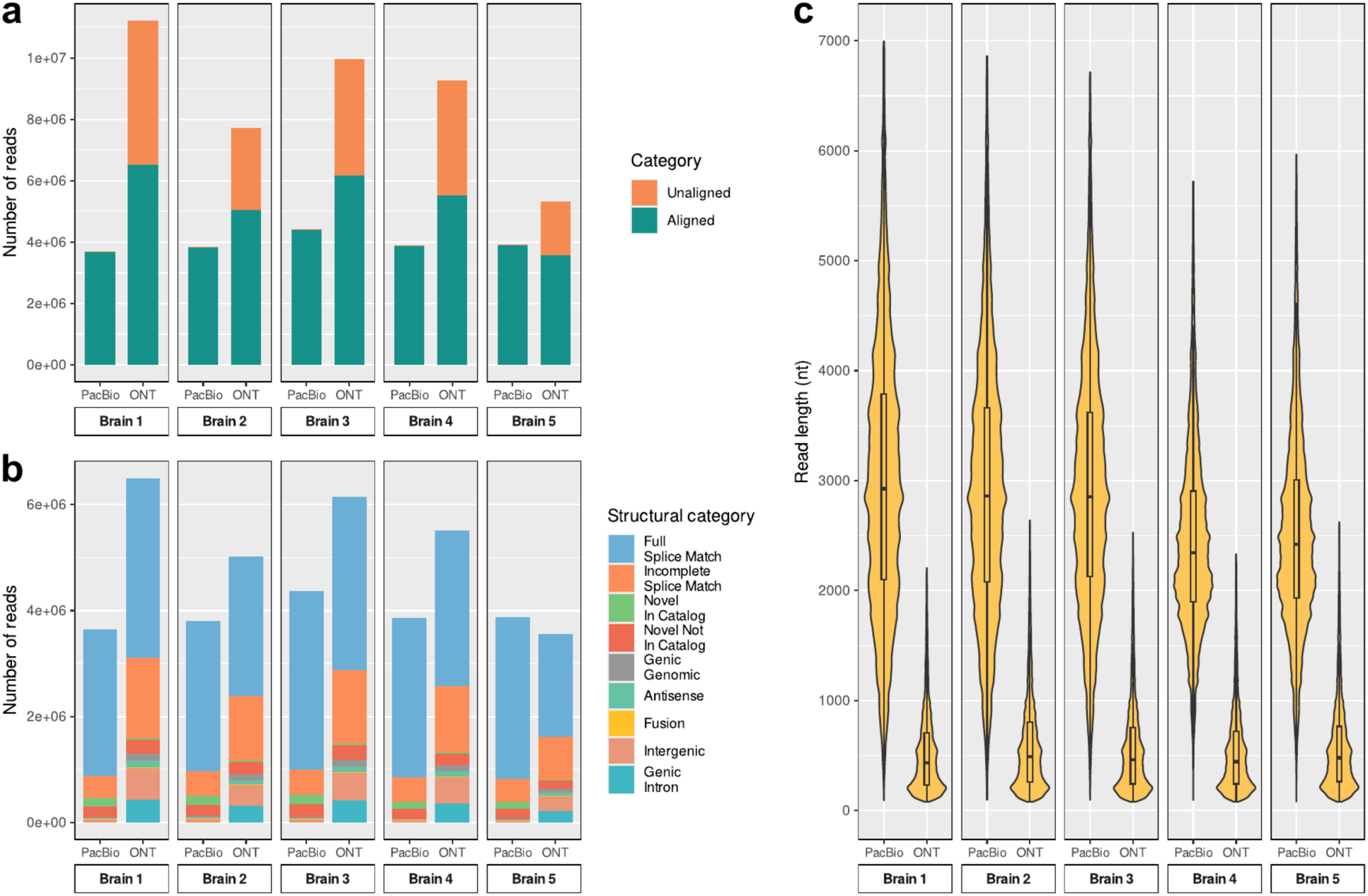
Read-level analysis on brain tissue. **a)** Number of aligned and unaligned reads per sample and technology on brain tissue. **b)** Structural categories of reads per sample and technology on brain tissue, according to analysis with SQANTI-Reads. **c)** Read-length distribution per sample and technology on brain tissue; Outliers were excluded by trimming each violin at the 99.5th percentile.

**Supplementary Figure 2:**
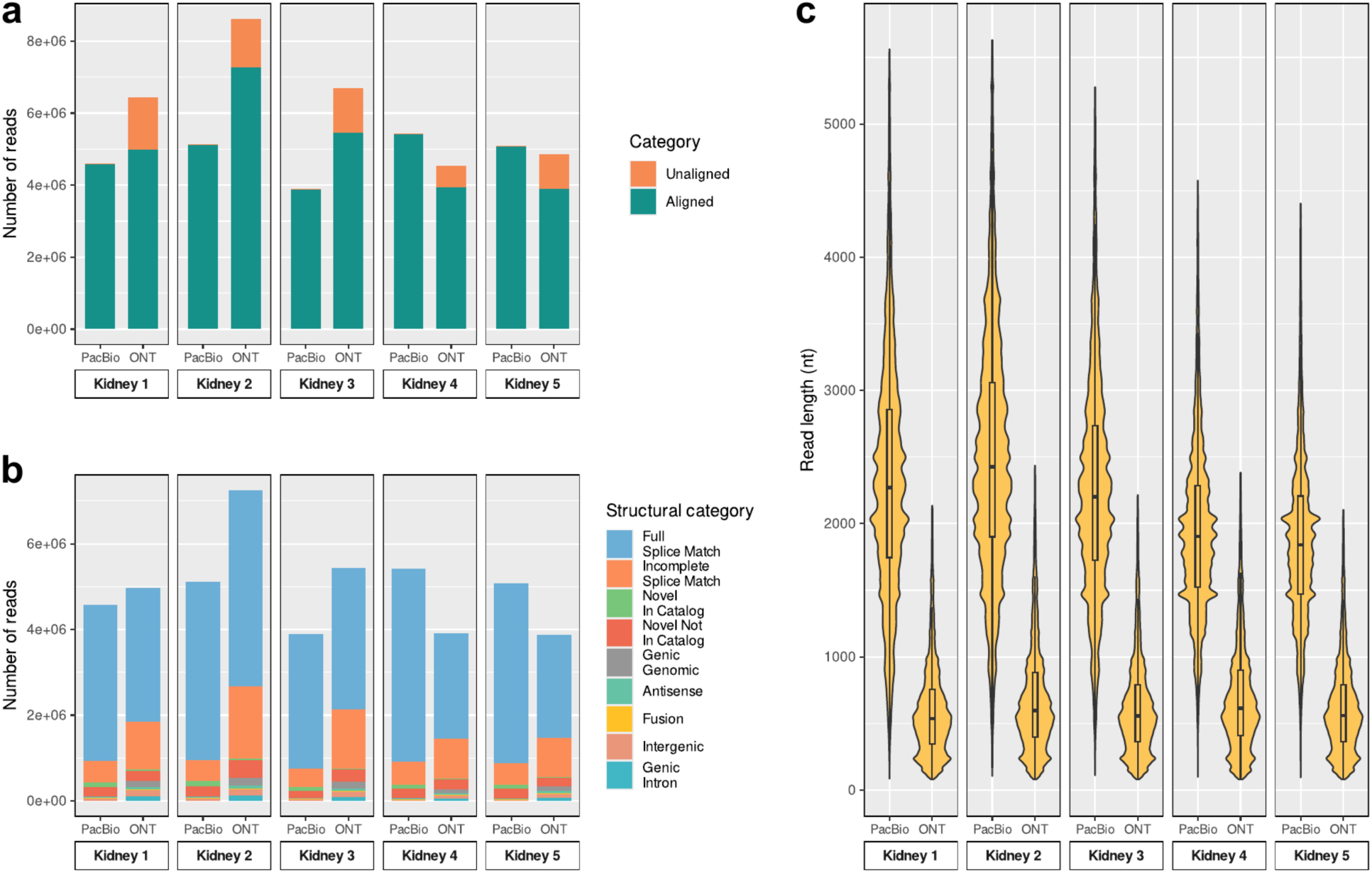
Read-level analysis on kidney tissue. **a)** Number of aligned and unaligned reads per sample and technology on kidney tissue. **b)** Structural categories of reads per sample and technology on kidney tissue, according to analysis with SQANTI-Reads. **c)** Read-length distribution per sample and technology on kidney tissue; Outliers were excluded by trimming each violin at the 99.5th percentile.

**Supplementary Figure 3:**
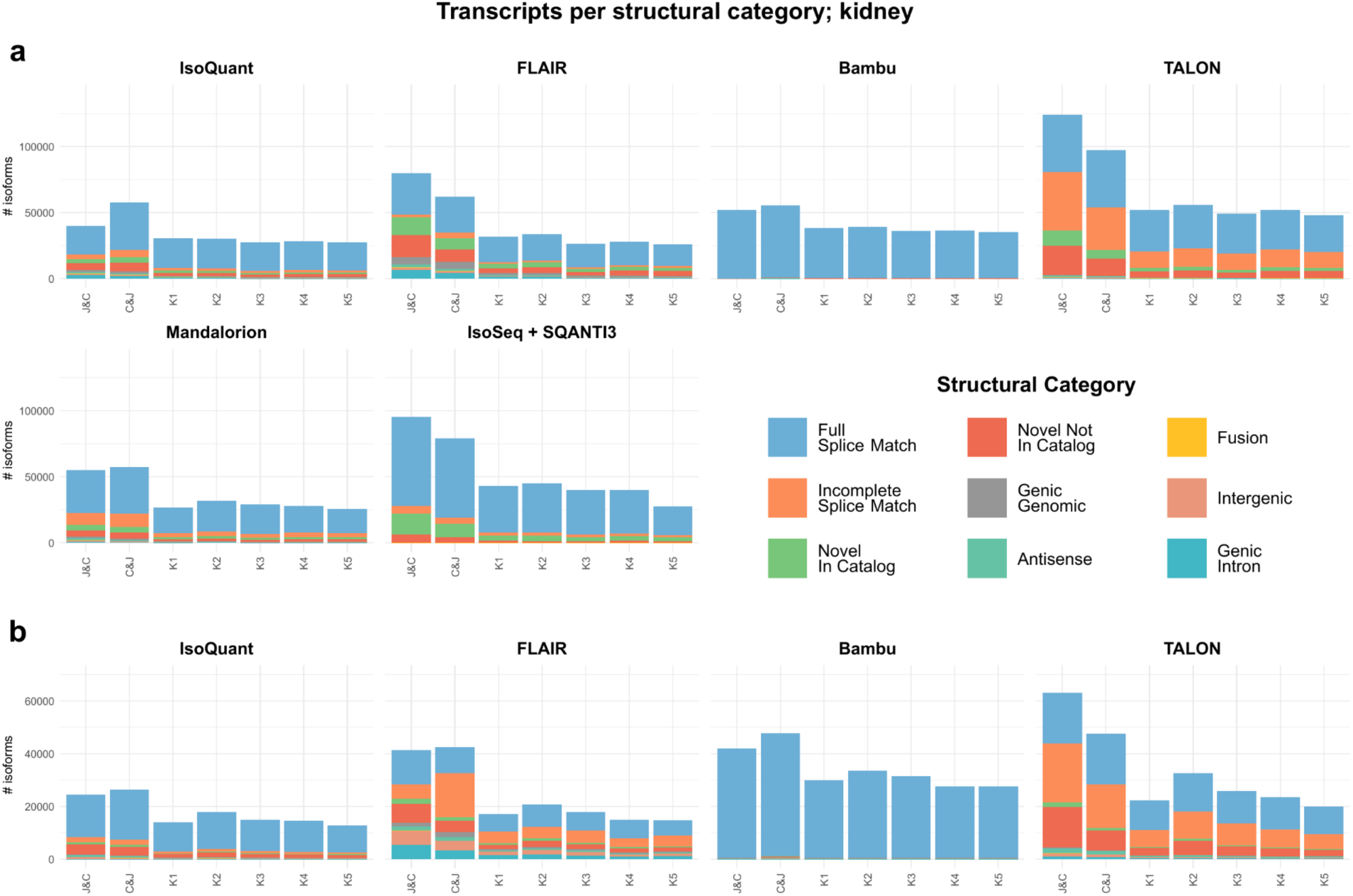
Transcripts per structural category in kidney tissue. Transcriptome diversity as measured by SQANTI3 structural categories across combination strategies and individual samples and across transcriptome reconstruction tools in **a)** PacBio and **b)** ONT data of mouse kidney tissue.

**Supplementary Figure 4:**
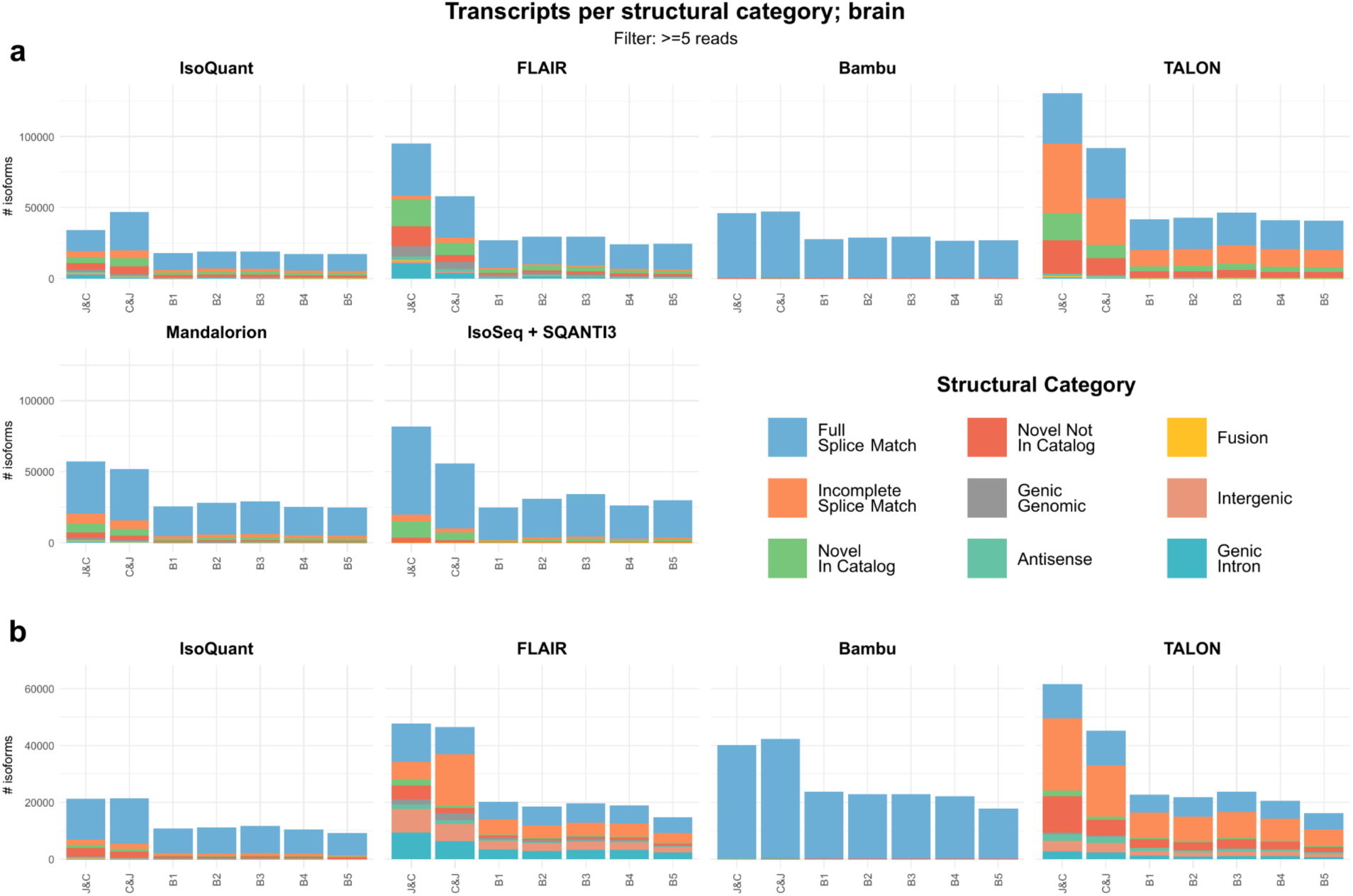
Transcripts with >= 5 reads per structural category in brain tissue. Transcriptome diversity as measured by SQANTI3 structural categories across combination strategies and individual samples and across transcriptome reconstruction tools in **a)** PacBio and **b)** ONT data of mouse brain tissue. Filtered to transcripts with a minimum expression of 5.

**Supplementary Figure 5:**
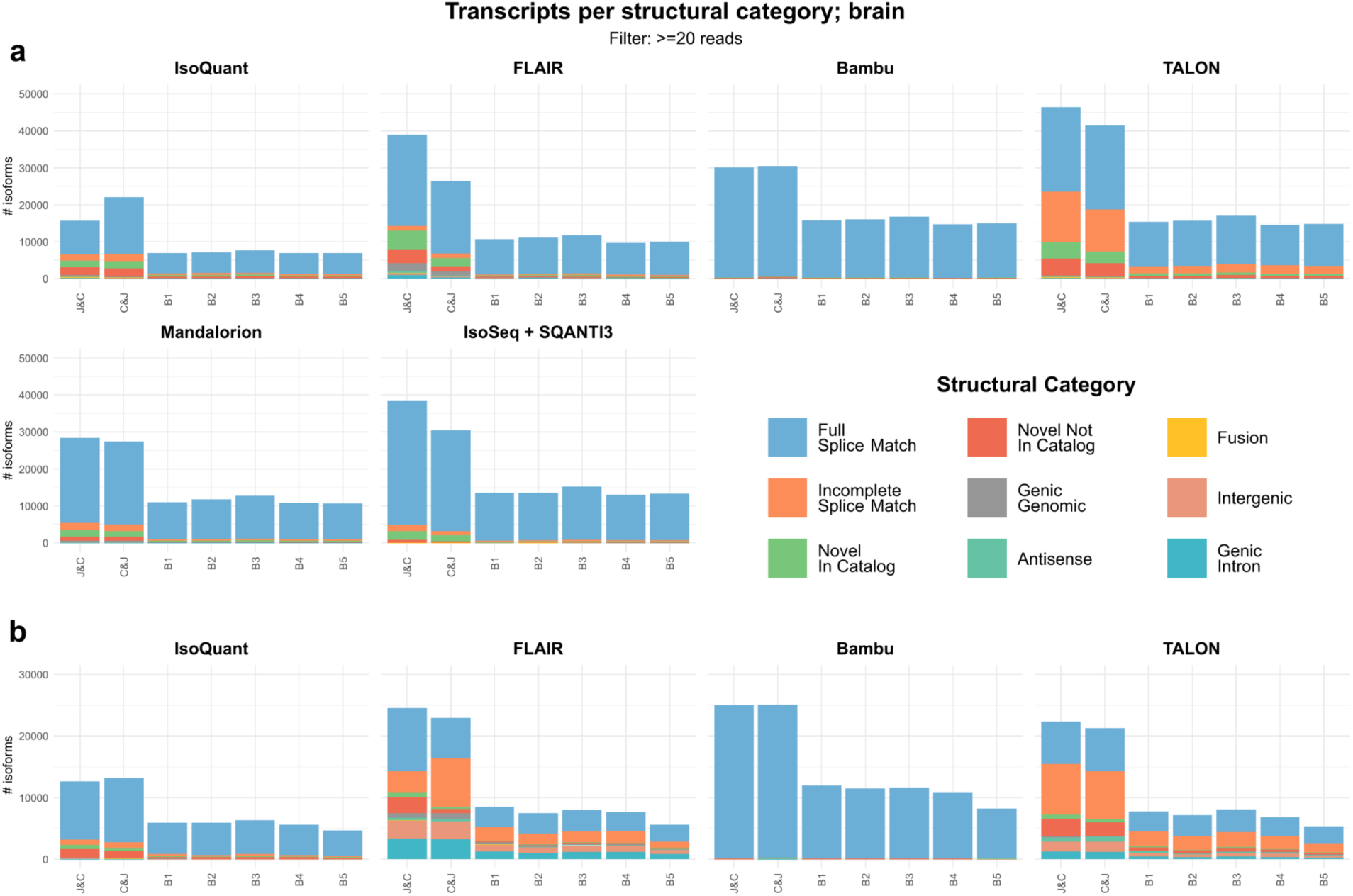
Transcripts with >= 20 reads per structural category in brain tissue. Transcriptome diversity as measured by SQANTI3 structural categories across combination strategies and individual samples and across transcriptome reconstruction tools in **a)** PacBio and **b)** ONT data of mouse brain tissue. Filtered to transcripts with a minimum expression of 10.

**Supplementary Figure 6:**
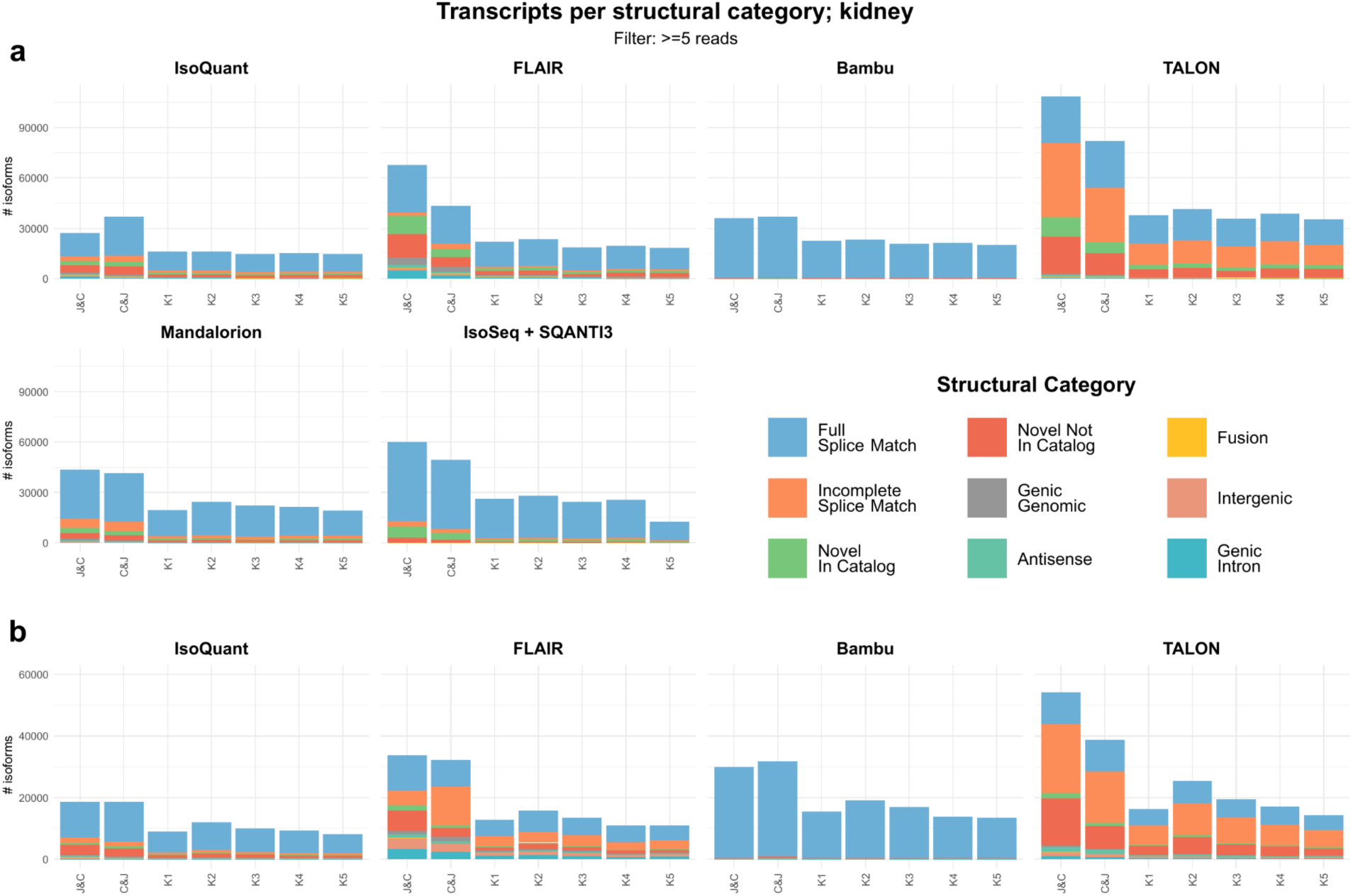
Transcripts with >= 5 reads per structural category in kidney tissue. Transcriptome diversity as measured by SQANTI3 structural categories across combination strategies and individual samples and across transcriptome reconstruction tools in **a)** PacBio and **b)** ONT data of mouse kidney tissue. Filtered to transcripts with a minimum expression of 5.

**Supplementary Figure 7:**
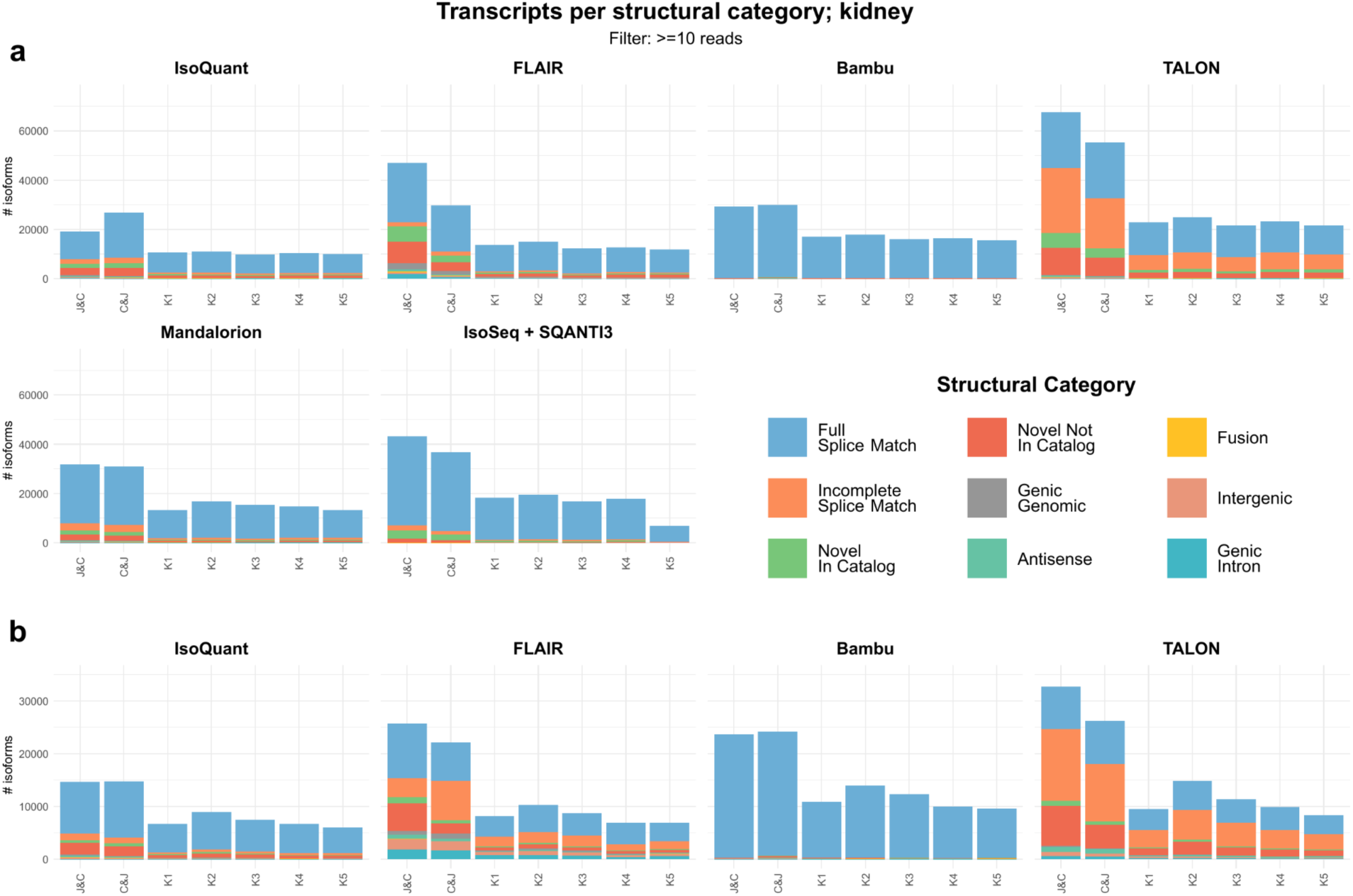
Transcripts with >= 10 reads per structural category in kidney tissue. Transcriptome diversity as measured by SQANTI3 structural categories across combination strategies and individual samples and across transcriptome reconstruction tools in **a)** PacBio and **b)** ONT data of mouse kidney tissue. Filtered to transcripts with a minimum expression of 10.

**Supplementary Figure 8:**
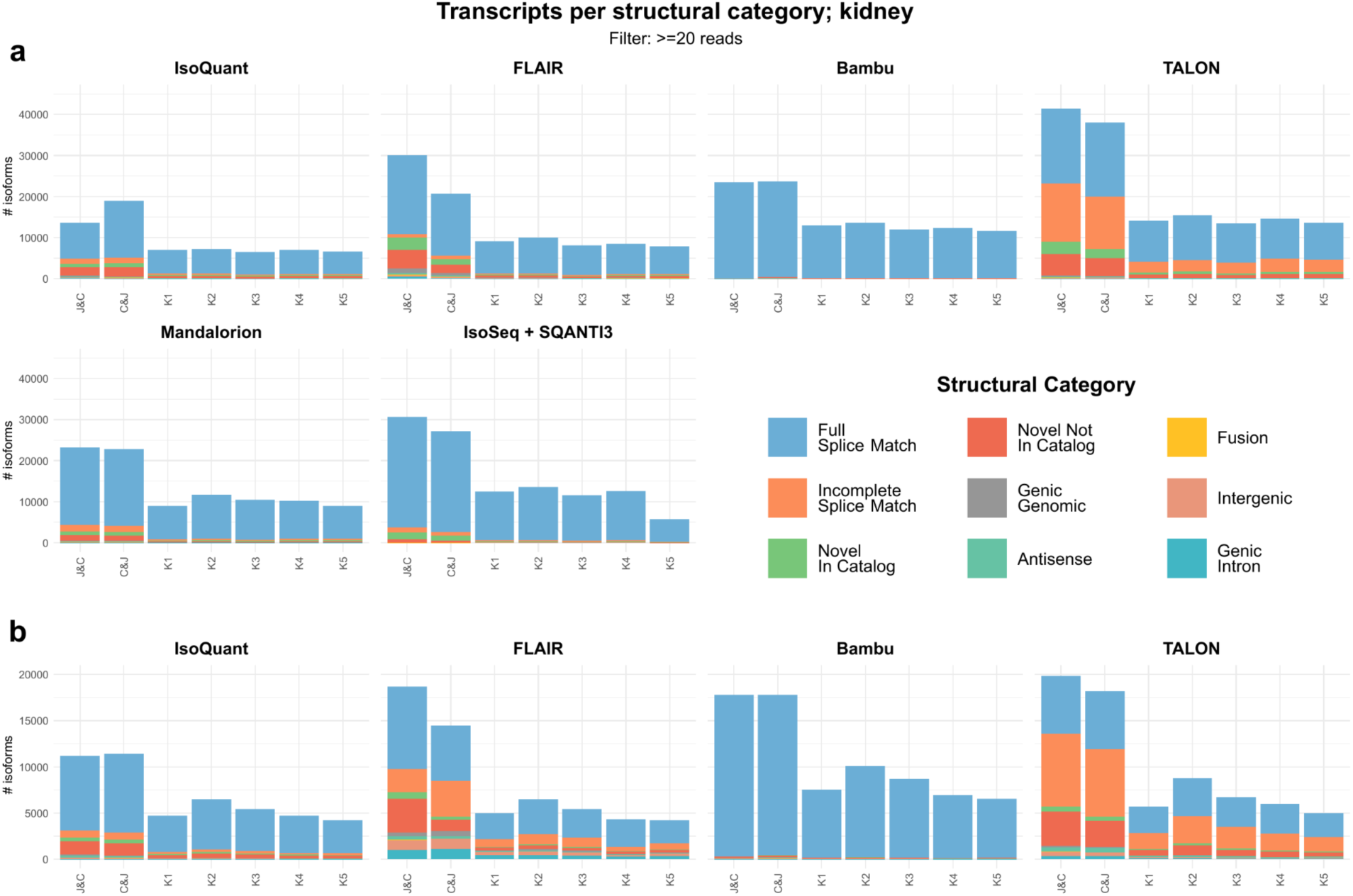
Transcripts with >= 20 reads per structural category in kidney tissue. Transcriptome diversity as measured by SQANTI3 structural categories across combination strategies and individual samples and across transcriptome reconstruction tools in **a)** PacBio and **b)** ONT data of mouse kidney tissue. Filtered to transcripts with a minimum expression of 20.

**Supplementary Figure 9:**
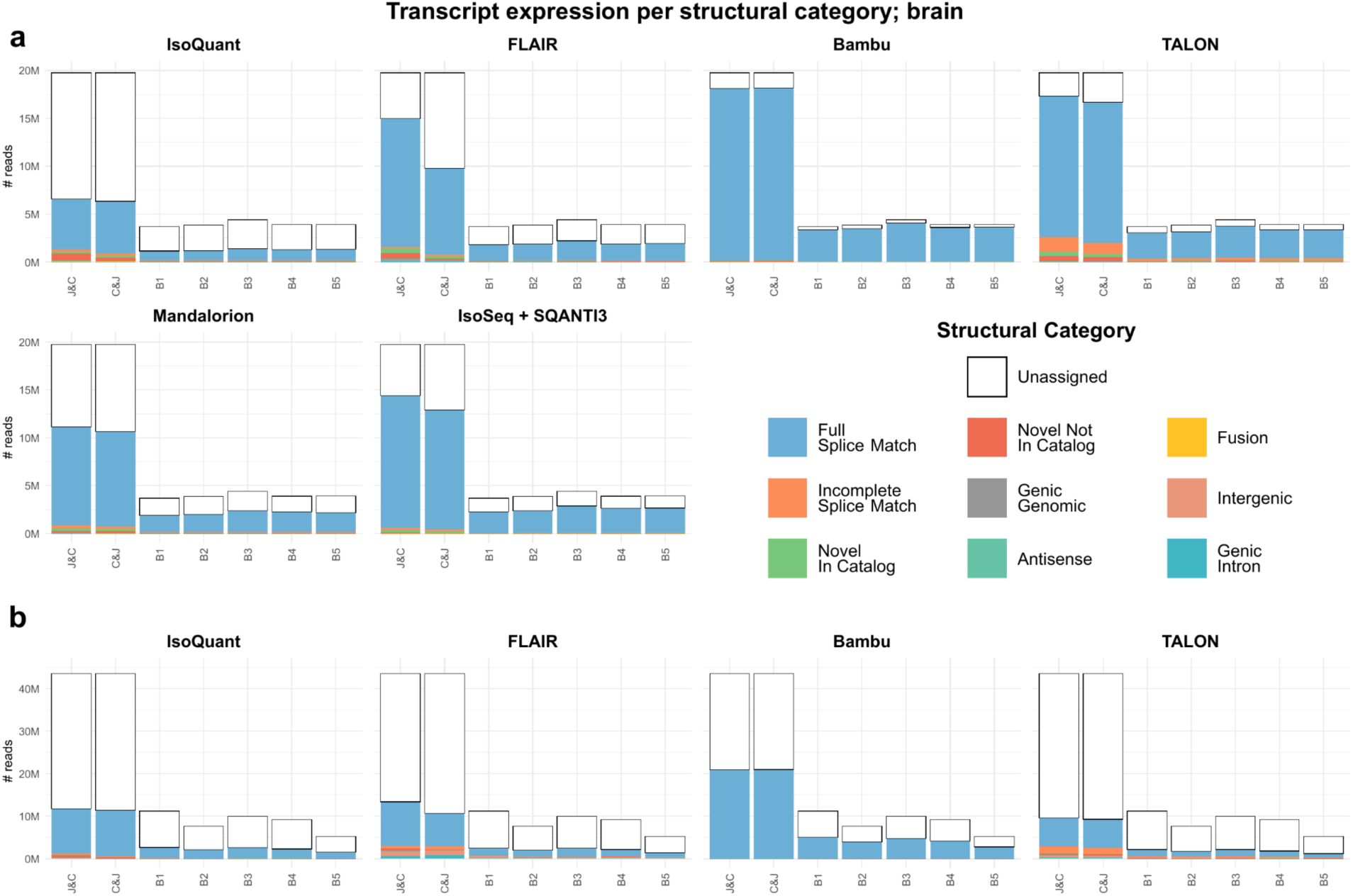
Reads per structural category of assigned transcript in brain tissue. Transcriptome diversity by number of reads as measured by SQANTI3 Quality Control’s structural categories of assigned transcripts across combination strategies and individual samples and across transcriptome reconstruction tools in **a)** PacBio and **b)** ONT data of mouse brain tissue. The “Unassigned” category includes reads that were either unmapped by minimap2, or not assigned to any isoform by the transcriptome reconstruction tool. Despite the potential for a large number of novel transcripts, the vast majority of reads were assigned to known (reference-annotated) transcripts, with novel transcripts generally being relatively lowly expressed. Interestingly, we observed strong differences in the number of unassigned reads across data types, with our ONT data exhibiting more unassigned reads. Furthermore, for some tools (FLAIR, TALON, IsoSeq with PacBio data; FLAIR with ONT data), J&C was capable of assigning more reads than C&J, aligning with the premise that pooled evidence enhances the ability to call lowly-expressed novel transcripts with greater confidence, thereby enabling more reads to be assigned.

**Supplementary Figure 10:**
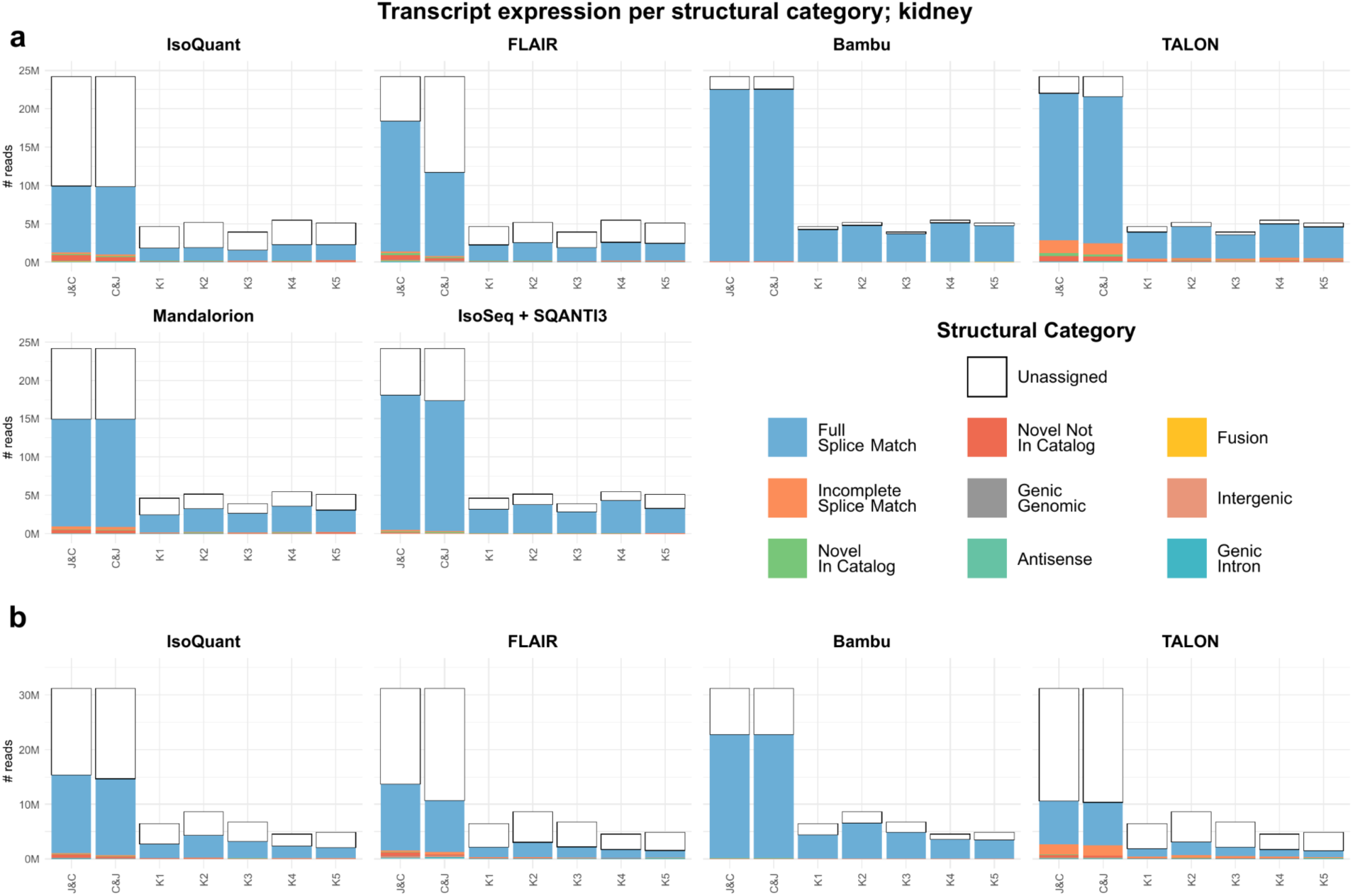
Unique Junction Chain overlap in kidney tissue. 10 biggest intersections of unique junction chains (UJCs) across combinations strategies and individual samples per transcriptome reconstruction tool for **a)** PacBio and **b)** ONT data of mouse kidney tissue.

**Supplementary Figure 11:**
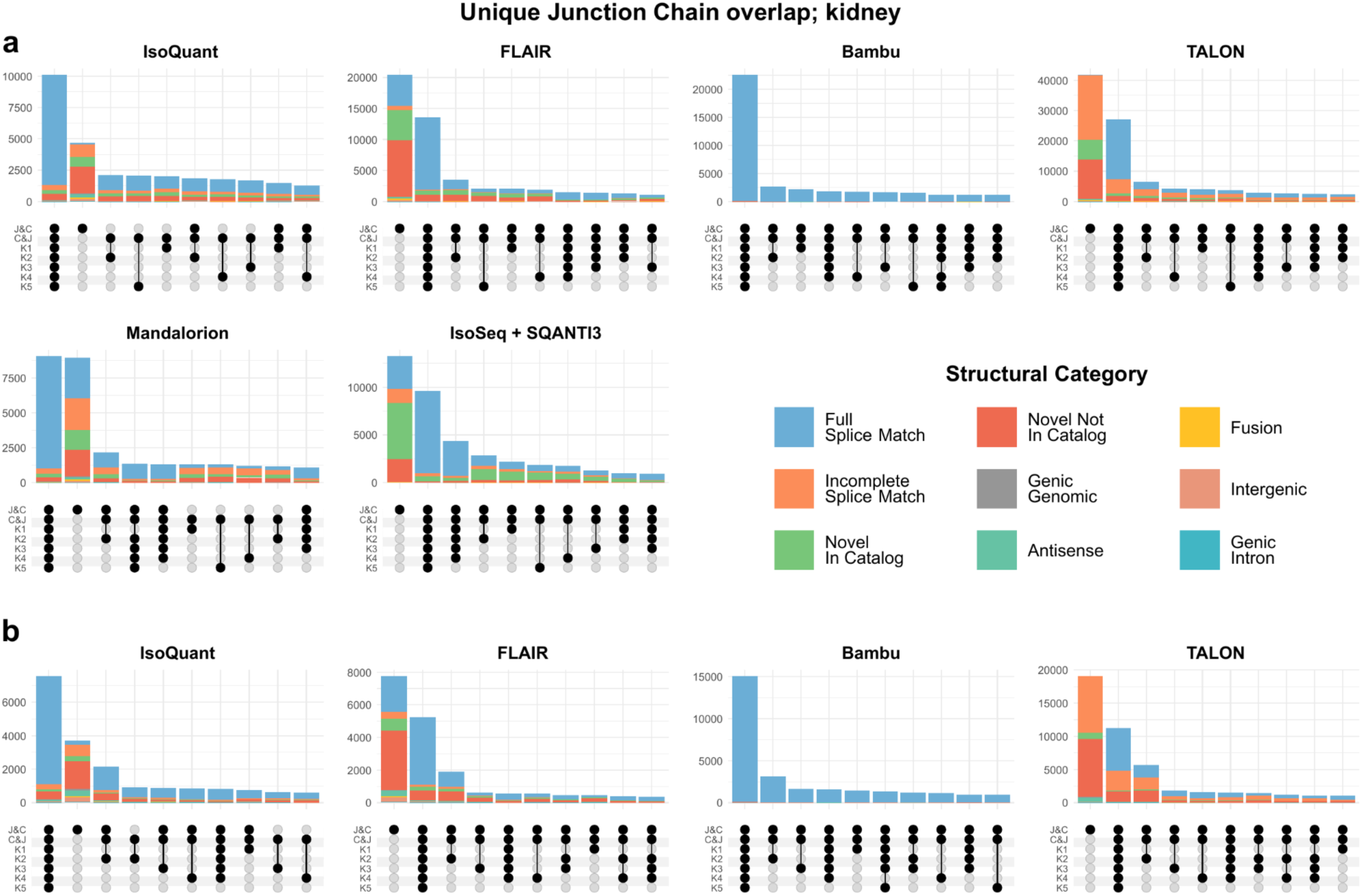
Unique Junction Chain overlap in kidney tissue. 10 biggest overlaps of unique junction chains (UJCs) across combinations strategies and individual samples per transcriptome reconstruction tool for **a)** PacBio and **b)** ONT data of mouse kidney tissue.

**Supplementary Figure 12:**
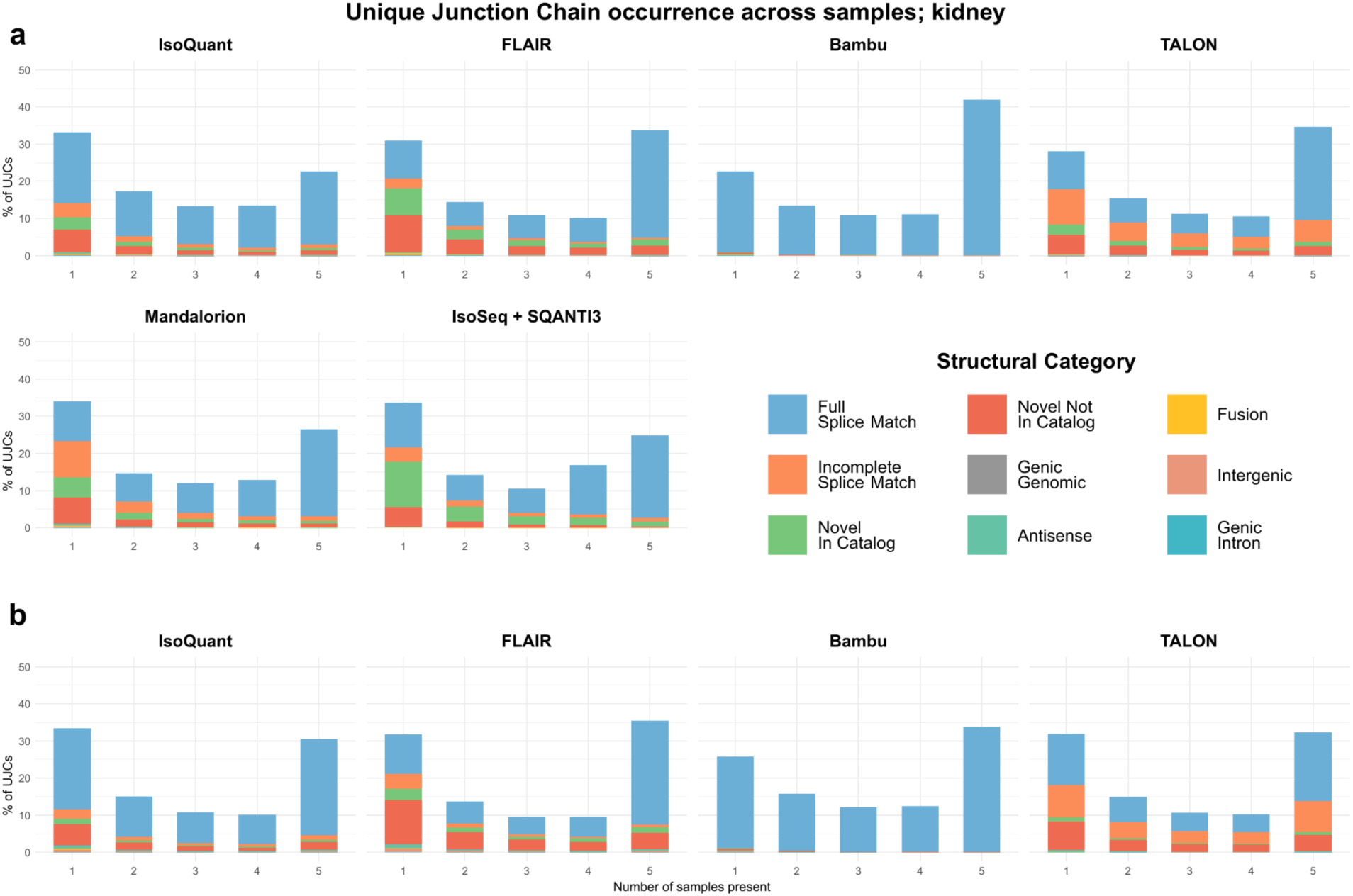
Distribution of unique junction chains (UJCs) across samples in kidney tissue. Distribution of UJC occurrence across number of samples by structural category per transcriptome reconstruction tool for **a)** PacBio and **b)** ONT data of mouse kidney tissue.

**Supplementary Figure 13:**
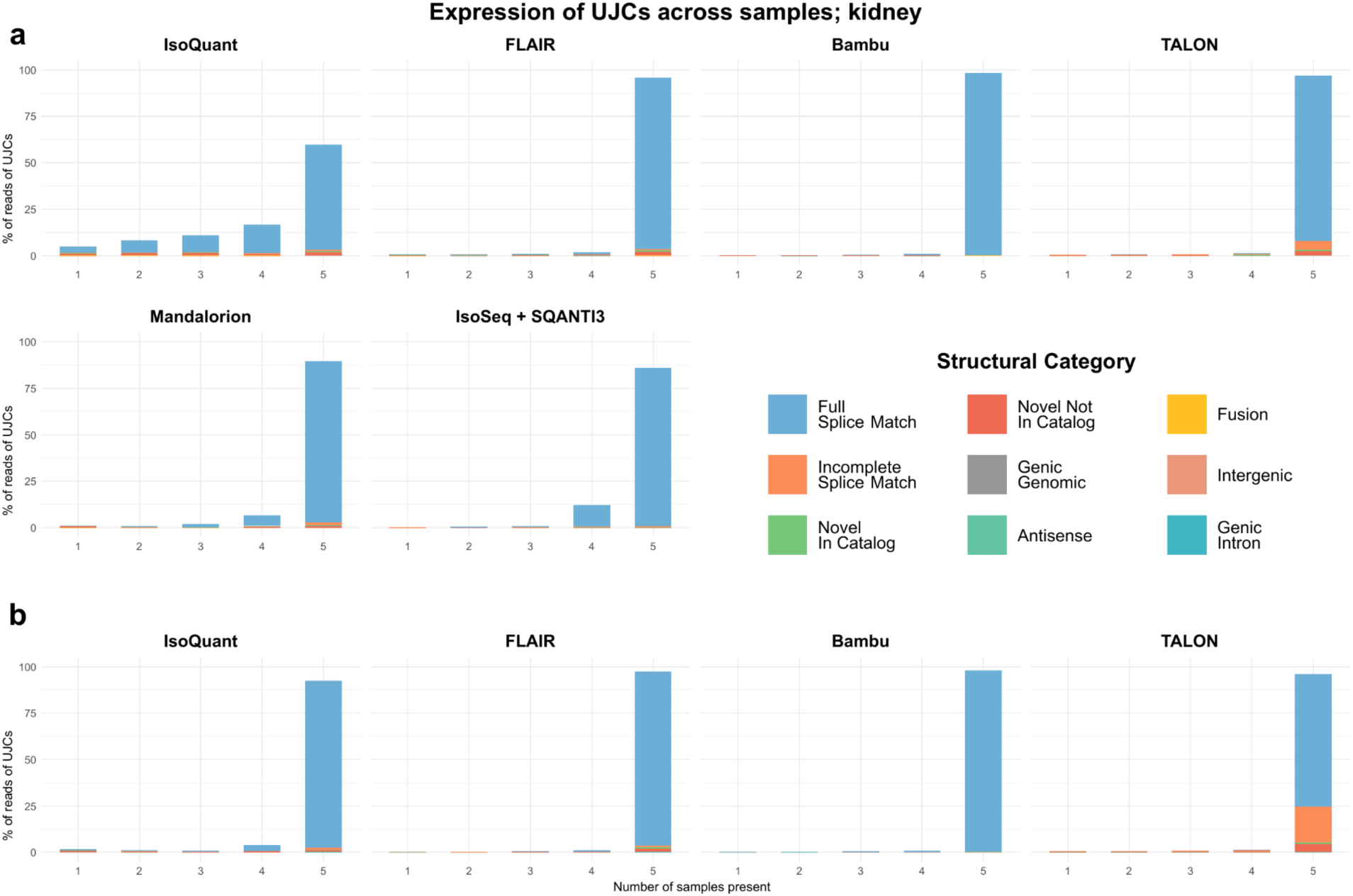
Distribution of expression of unique junction chains (UJCs) across samples in kidney tissue. Distribution of expression of UJC occurrence across number of samples by structural category per transcriptome reconstruction tool for **a)** PacBio and **b)** ONT data of mouse kidney tissue.

**Supplementary Figure 14:**
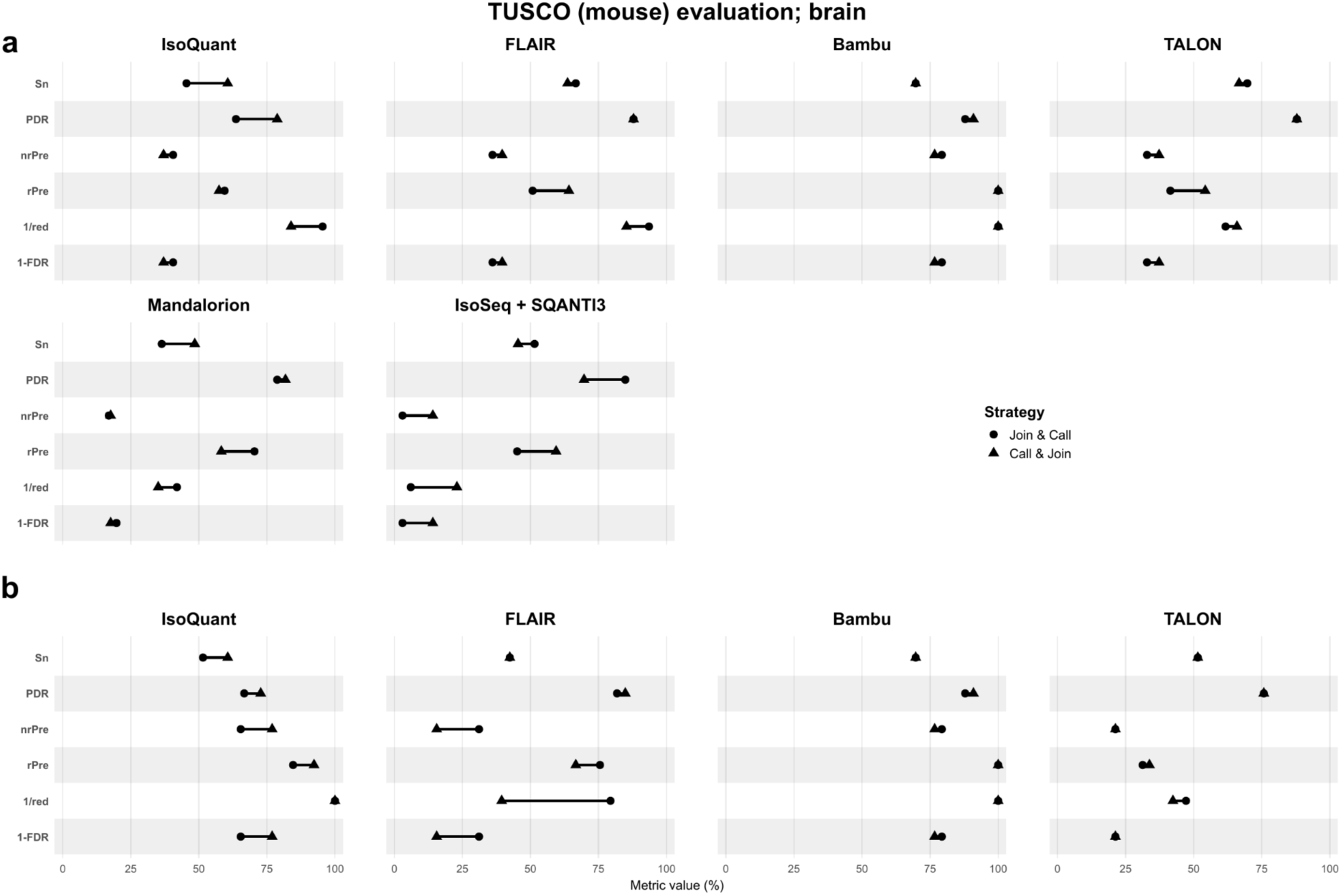
TUSCO (mouse) evaluation in brain tissue. Sensitivity (Sn), positive detection rate (PDR), non-redundant precision (nrPre), redundant precision (rPre), redundancy (red) and false discovery rate (FDR) for TUSCO (mouse) analysis comparing the combination strategies per transcriptome reconstruction tool for **a)** PacBio and **b)** ONT data of mouse brain tissue.

**Supplementary Figure 15:**
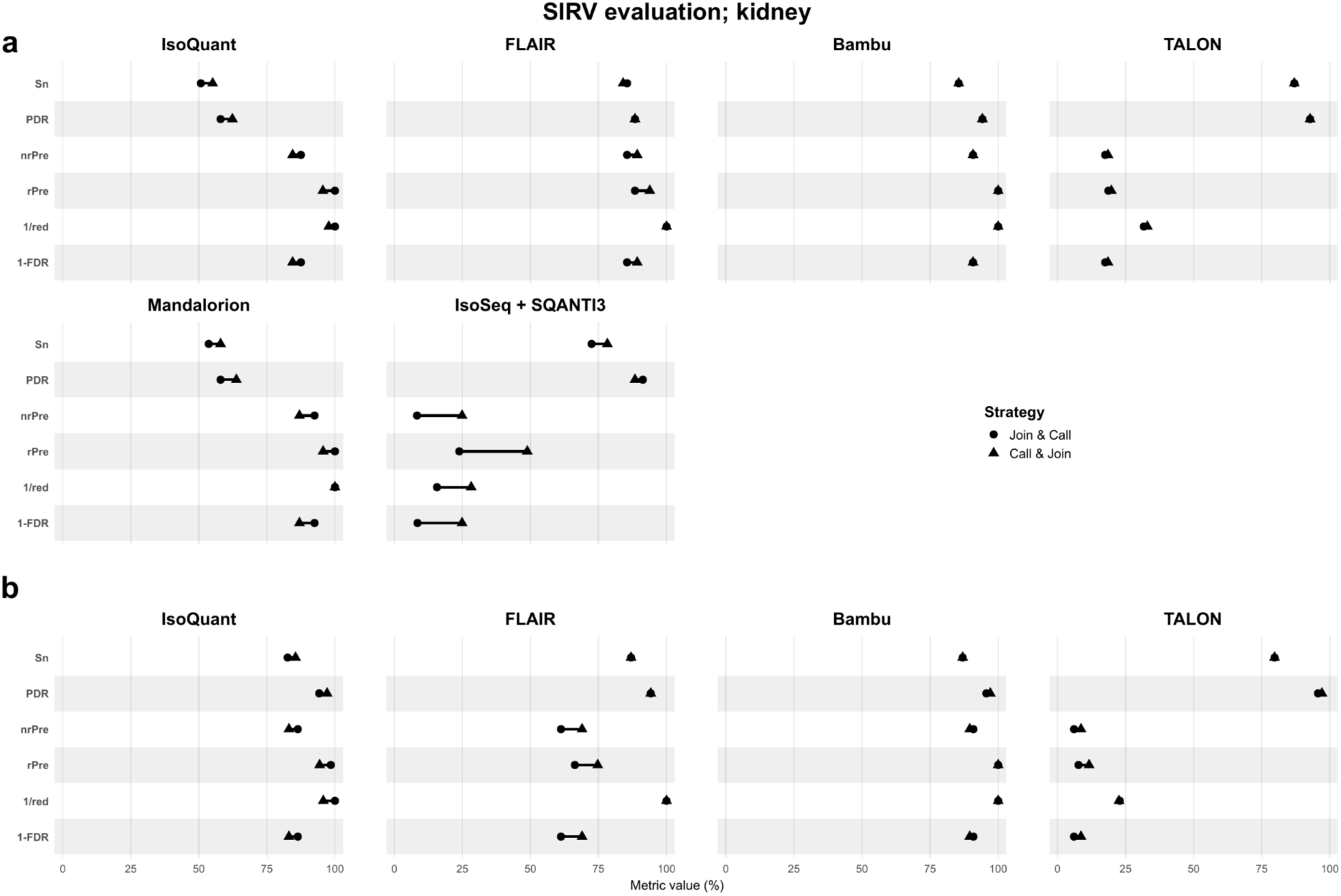
SIRV evaluation in kidney tissue. Sensitivity (Sn), positive detection rate (PDR), non-redundant precision (nrPre), redundant precision (rPre), redundancy (red) and false discovery rate (FDR) for TUSCO (mouse) and SIRV analyses comparing the combination strategies per transcriptome reconstruction tool for **a)** PacBio and **b)** ONT data of mouse kidney tissue.

**Supplementary Figure 16:**
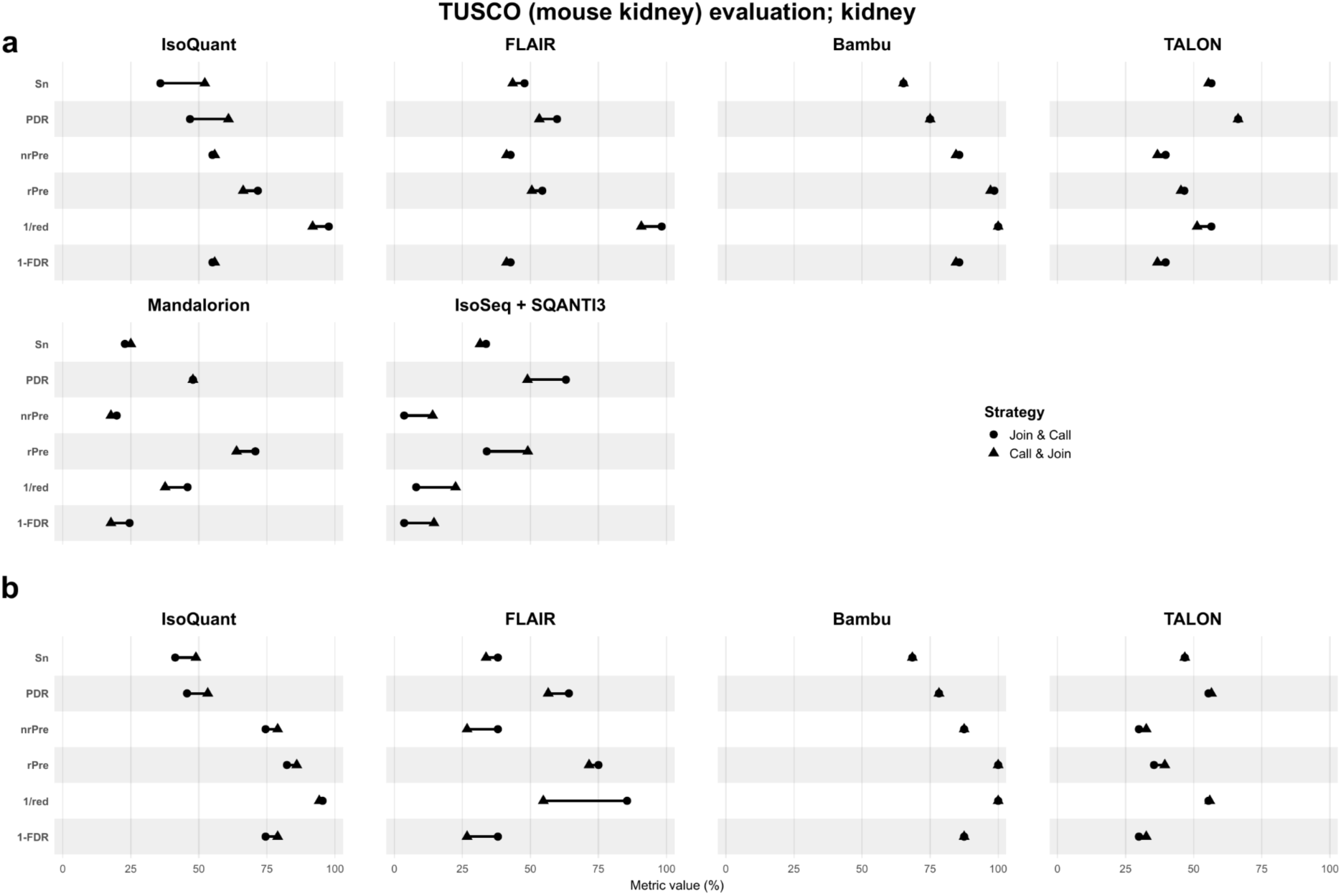
TUSCO (mouse kidney) evaluation in kidney tissue. Sensitivity (Sn), positive detection rate (PDR), non-redundant precision (nrPre), redundant precision (rPre), redundancy (red) and false discovery rate (FDR) for TUSCO (mouse kidney) analysis comparing the combination strategies per transcriptome reconstruction tool for **a)** PacBio and **b)** ONT data of mouse kidney tissue.

**Supplementary Figure 17:**
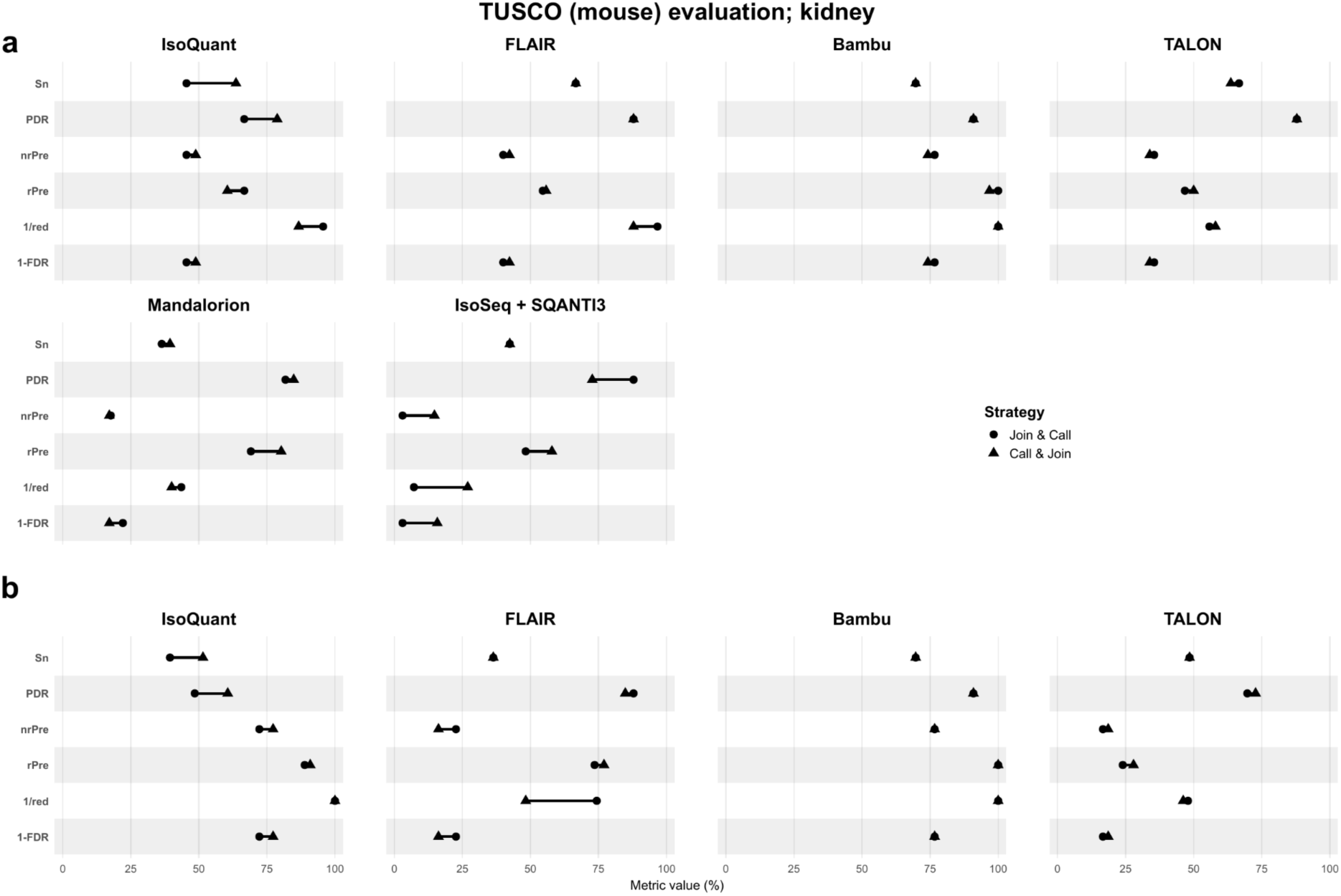
TUSCO (mouse) evaluation in kidney tissue. Sensitivity (Sn), positive detection rate (PDR), non-redundant precision (nrPre), redundant precision (rPre), redundancy (red) and false discovery rate (FDR) for TUSCO (mouse) analysis comparing the combination strategies per transcriptome reconstruction tool for **a)** PacBio and **b)** ONT data of mouse kidney tissue.

**Supplementary Figure 18:**
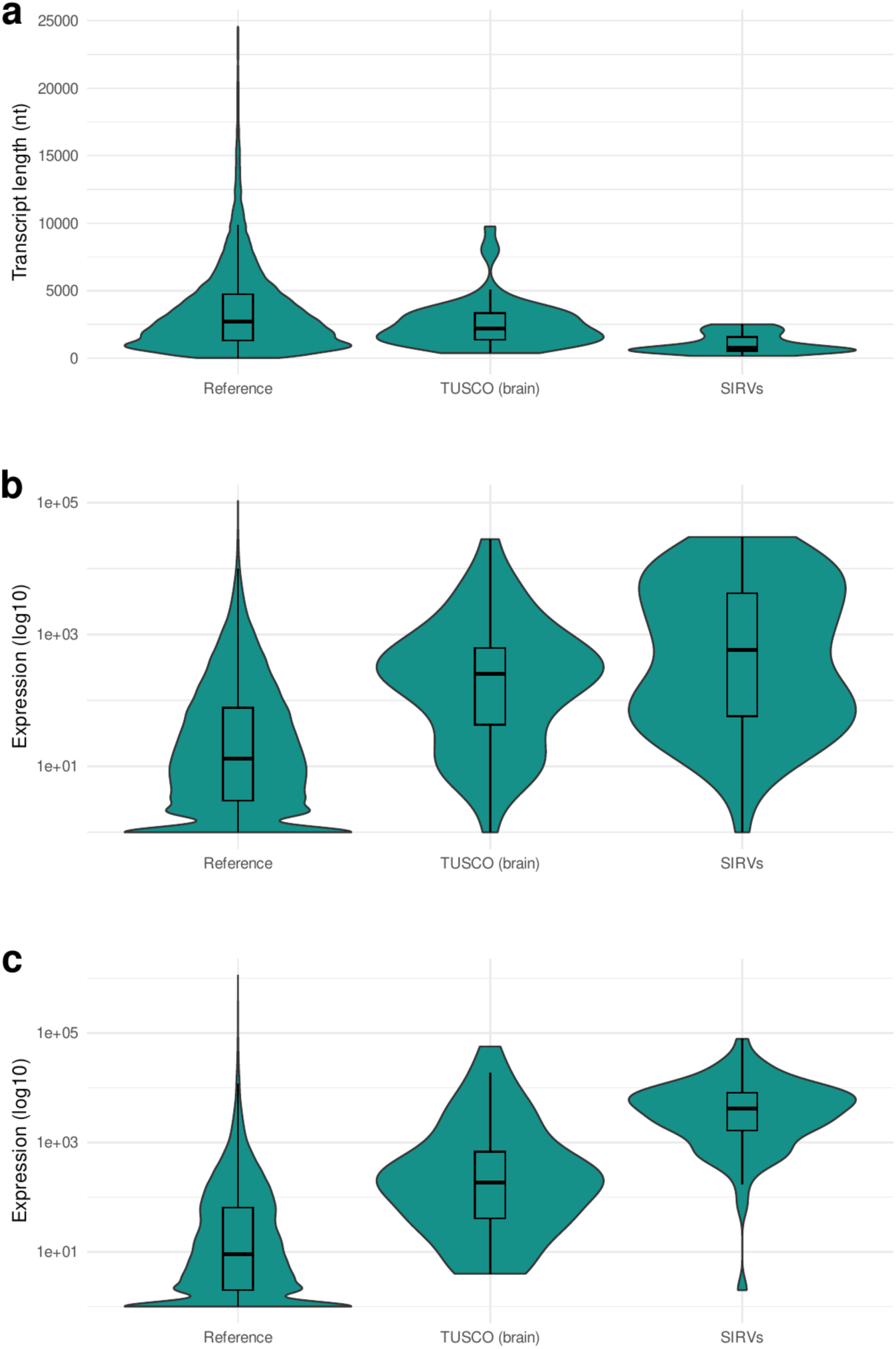
Comparison of reference, TUSCO, and SIRV transcripts in brain tissue. **a)** Length of reference, TUSCO (mouse brain), and SIRV transcripts. Reference transcripts were trimmed at the 99.5th percentile to exclude outliers. **b)** Expression of reference, TUSCO (mouse brain), and SIRV transcripts in PacBio data based on SQANTI-Reads analysis. **c)** Expression of reference, TUSCO (mouse brain), and SIRV transcripts in ONT data based on SQANTI-Reads analysis.

**Supplementary Figure 19:**
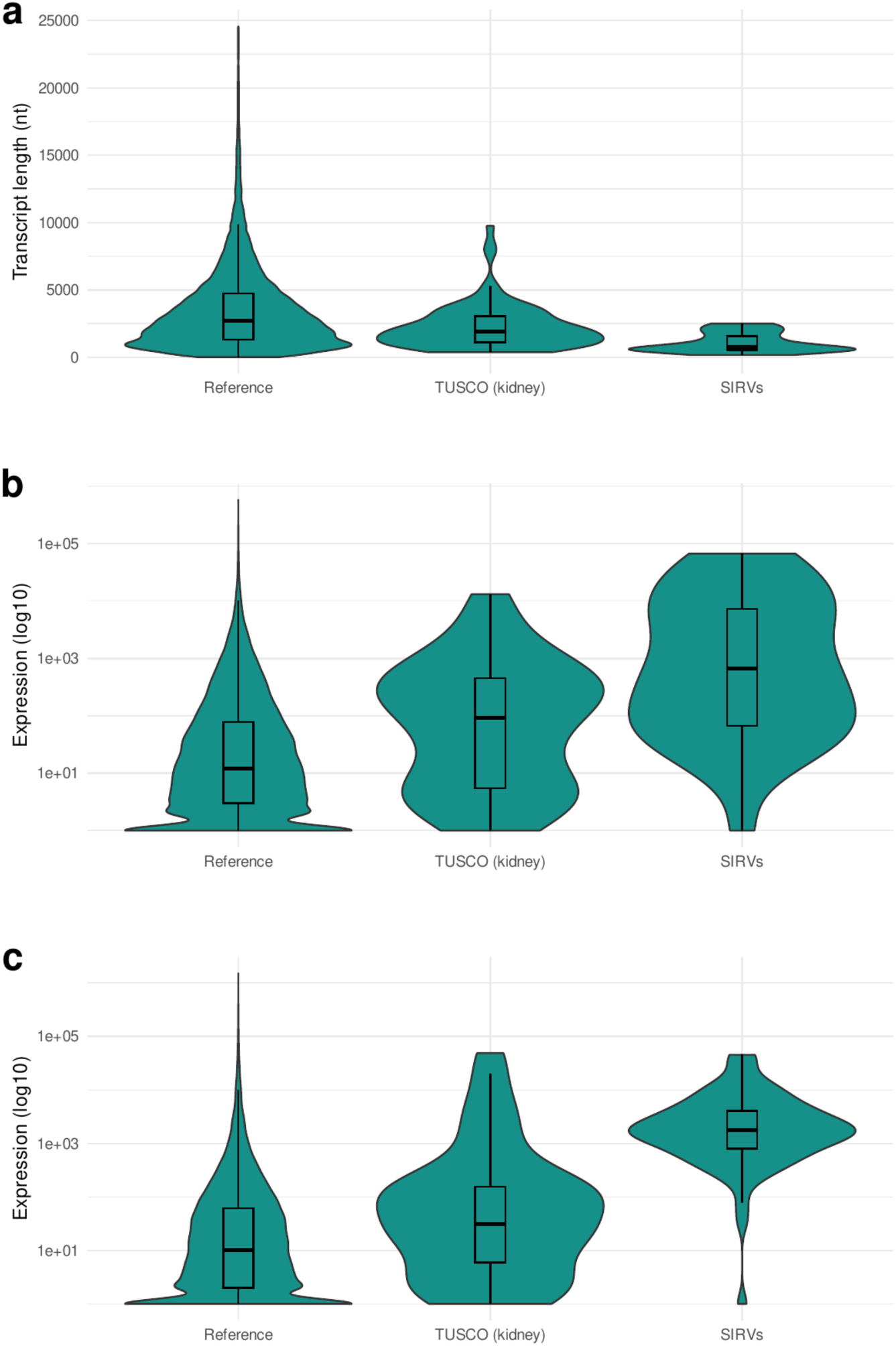
Comparison of reference, TUSCO, and SIRV transcripts in kidney tissue. **a)** Length of reference, TUSCO (mouse kidney), and SIRV transcripts. Reference transcripts were trimmed at the 99.5th percentile to exclude outliers. **b)** Expression of reference, TUSCO (mouse kidney), and SIRV transcripts in PacBio data based on SQANTI-Reads analysis. **c)** Expression of reference, TUSCO (mouse kidney), and SIRV transcripts in ONT data based on SQANTI-Reads analysis.

**Supplementary Figure 20:**
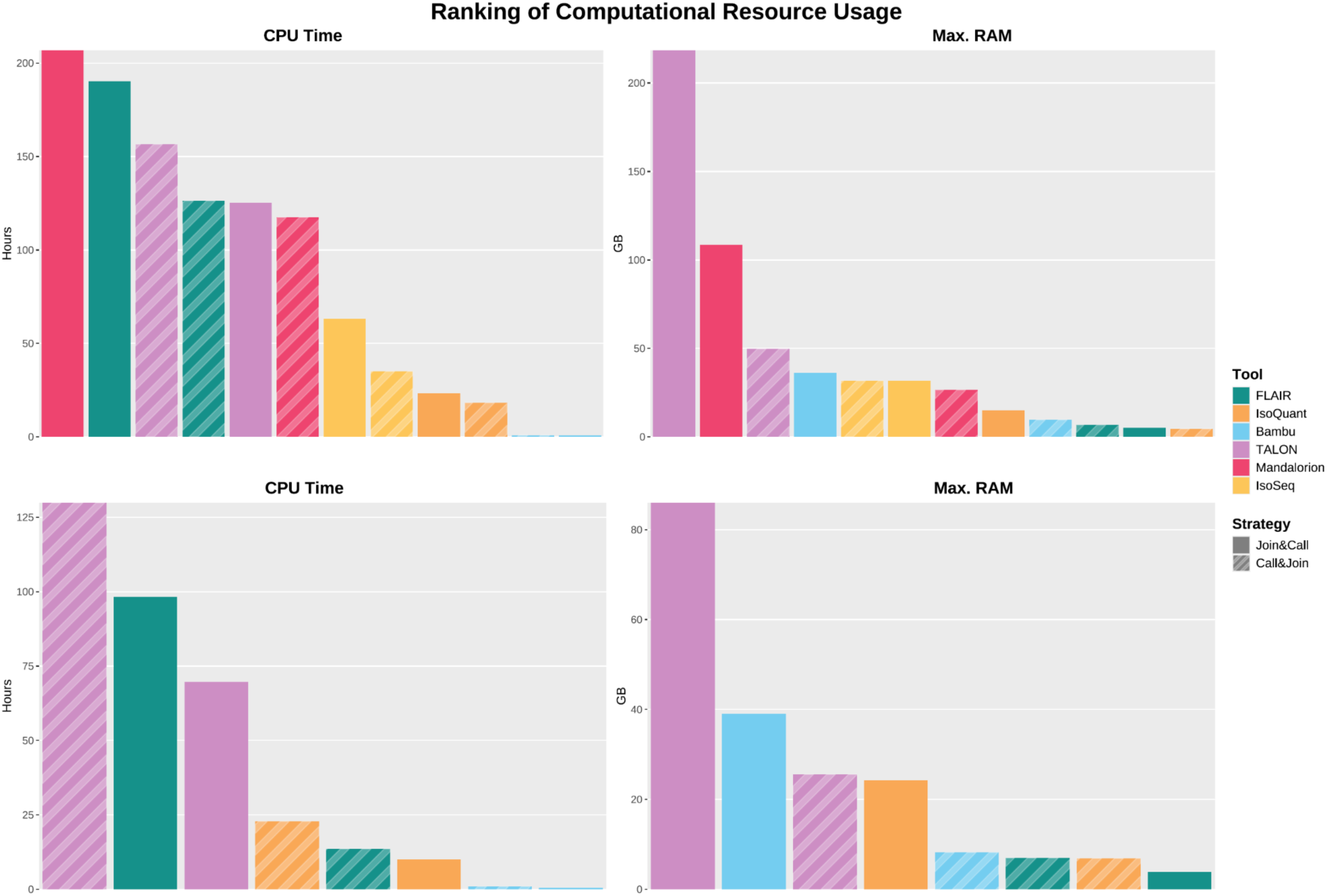
Ranking of CPU Time and Maximum RAM Usage in kidney tissue. Ranking of transcriptome reconstruction tools and combination strategies by their required CPU Time in hours and maximum memory usage in gigabytes in **a)** PacBio and **b)** ONT data of mouse kidney tissue. For C&J, the CPU time measurements are the sum of the individual processes for each sample plus the process for TAMA Merge, while the maximum memory is the maximum of the individual processes. For J&C, wherever processes were split (e.g. by chromosome or for identification and quantification), the measurements were aggregated in the same way.

**Supplementary Table A:**
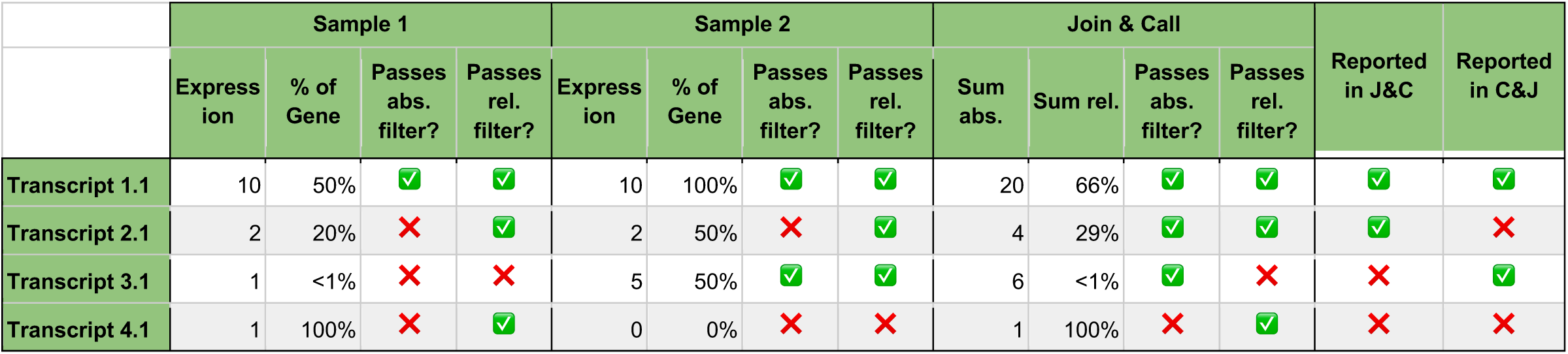
Impact of relative expression filters on transcript detection in combination strategies. This table demonstrates 4 hypothetical transcripts belonging to 4 different genes with different expression profiles across 2 samples. An absolute filter of >=3 reads and a relative filter of >=2% of gene expression are applied. Transcript 1.1 is comparatively highly expressed, surpassing both filters in both samples, therefore also being reported in both strategies. Transcript 2.1 is lowly expressed in both samples. It is not detected when samples are analyzed individually in C&J, but can be detected in J&C by combining evidence from both samples. Transcript 3.1 fails both filters in one sample, but passes both in the other. As it is detected in sample 2, when combining the resulting transcript models, C&J reports Transcript 3.1. However, when combining reads from both samples, it fails to pass the relative threshold of 2% due to high expression of other transcripts of the same gene in sample 1 and is therefore not reported in J&C. Transcript 4.1 is generally lowly expressed and, even when combining evidence, does not surpass the absolute filter. It therefore goes unreported in both strategies.

**Supplementary Table B:** Computational Resources. Computational resources (elapsed and CPU time, maximum memory) used by each combination of data type, tissue, tool, and strategy. For C&J, the elapsed and CPU time measurements are the sum of the individual processes for each sample plus the process for TAMA Merge, while the maximum memory is the maximum of the individual processes. For J&C, wherever processes were split (e.g. by chromosome or for identification and quantification), the measurements were aggregated in the same way.

| Data Type | Tissue | Tool | Strategy | Elapsed Time | CPU Time | Max. RAM |
| --- | --- | --- | --- | --- | --- | --- |
| ONT | brain | IsoQuant | Call&Join | 0-05:40:51 | 0-16:10:55 | 6.8GB |
| ONT | kidney | IsoQuant | Call&Join | 0-04:42:31 | 0-22:46:38 | 6.9GB |
| ONT | brain | IsoQuant | Join&Call | 0-08:33:23 | 0-15:17:16 | 22.0GB |
| ONT | kidney | IsoQuant | Join&Call | 0-08:01:09 | 0-10:03:01 | 24.2GB |
| ONT | brain | FLAIR | Call&Join | 0-01:55:23 | 0-15:53:35 | 7.3GB |
| ONT | kidney | FLAIR | Call&Join | 0-01:37:23 | 0-13:36:13 | 7.0GB |
| ONT | brain | FLAIR | Join&Call | 1-16:53:25 | 3-16:24:40 | 4.2GB |
| ONT | kidney | FLAIR | Join&Call | 1-19:49:07 | 4-02:16:41 | 3.9GB |
| ONT | brain | bambu | Call&Join | 0-00:49:28 | 0-00:52:47 | 10.5GB |
| ONT | kidney | bambu | Call&Join | 0-00:55:46 | 0-00:57:19 | 8.2GB |
| ONT | brain | bambu | Join&Call | 0-00:32:19 | 0-00:32:52 | 38.3GB |
| ONT | kidney | bambu | Join&Call | 0-00:30:13 | 0-00:30:57 | 39.0GB |
| ONT | brain | TALON | Call&Join | 0-11:53:39 | 3-21:17:10 | 20.1GB |
| ONT | kidney | TALON | Call&Join | 0-12:08:05 | 5-09:53:04 | 25.6GB |
| ONT | brain | TALON | Join&Call | 0-06:49:09 | 2-05:05:07 | 85.3GB |
| ONT | kidney | TALON | Join&Call | 0-07:04:08 | 2-21:45:18 | 86.0GB |
| PacBio | brain | IsoQuant | Call&Join | 0-03:01:21 | 0-19:43:37 | 3.2GB |
| PacBio | kidney | IsoQuant | Call&Join | 0-02:26:57 | 0-18:15:27 | 4.6GB |
| PacBio | brain | IsoQuant | Join&Call | 0-06:00:10 | 0-22:48:34 | 7.9GB |
| PacBio | kidney | IsoQuant | Join&Call | 0-05:10:34 | 0-23:18:43 | 15.1GB |
| PacBio | brain | FLAIR | Call&Join | 0-09:55:51 | 5-02:15:41 | 6.9GB |
| PacBio | kidney | FLAIR | Call&Join | 0-10:05:46 | 5-06:18:27 | 6.7GB |
| PacBio | brain | FLAIR | Join&Call | 4-20:16:13 | 10-12:59:14 | 6.2GB |
| PacBio | kidney | FLAIR | Join&Call | 2-23:11:21 | 7-22:22:57 | 5.0GB |
| PacBio | brain | bambu | Call&Join | 0-01:13:45 | 0-01:16:55 | 9.9GB |
| PacBio | kidney | bambu | Call&Join | 0-00:54:50 | 0-00:57:31 | 9.6GB |
| PacBio | brain | bambu | Join&Call | 0-00:54:49 | 0-00:55:59 | 32.9GB |
| PacBio | kidney | bambu | Join&Call | 0-00:46:35 | 0-00:47:53 | 36.2GB |
| PacBio | brain | TALON | Call&Join | 0-12:07:43 | 6-09:06:14 | 48.5GB |
| PacBio | kidney | TALON | Call&Join | 0-12:42:22 | 6-12:30:43 | 49.7GB |
| PacBio<br>PacBio | brain | TALON | Join&Call | 0-13:15:46 | 6-18:55:59 | 206.4GB |
| PacBio | kidney | TALON | Join&Call | 0-10:26:35 | 5-05:20:24 | 218.5GB |
| PacBio | brain | Mandalorion | Call&Join | 16-16:38:41 | 5-19:41:07 | 26.2GB |
| PacBio | kidney | Mandalorion | Call&Join | 18-01:04:51 | 4-21:31:20 | 26.5GB |
| PacBio | brain | Mandalorion | Join&Call | 1-03:30:01 | 10-06:11:09 | 111.4GB |
| PacBio | kidney | Mandalorion | Join&Call | 0-20:01:25 | 8-14:50:26 | 108.7GB |
| IsoSeq | brain | IsoSeq | Call&Join | 0-05:32:58 | 2-16:42:15 | 29.1GB |
| PacBio | kidney | IsoSeq | Call&Join | 0-03:11:12 | 1-11:03:04 | 31.7GB |
| PacBio | brain | IsoSeq | Join&Call | 0-07:18:45 | 3-12:01:42 | 31.0GB |
| PacBio | kidney | IsoSeq | Join&Call | 0-09:15:09 | 2-15:13:17 | 31.6GB |

## References

1. Monzó, C., Liu, T. & Conesa, A. Transcriptomics in the era of long-read sequencing. Nat Rev Genet 26, 681–701 (2025).

2. Tilgner, H. et al. Comprehensive transcriptome analysis using synthetic long-read sequencing reveals molecular co-association of distant splicing events. Nat Biotechnol 33, 736–742 (2015).

3. Glinos, D. A. et al. Transcriptome variation in human tissues revealed by long-read sequencing. Nature 608, 353–359 (2022).

4. Monzó, C., Frankish, A. & Conesa, A. Notable challenges posed by long-read sequencing for the study of transcriptional diversity and genome annotation. Genome Res 35, 583–592 (2025).

5. Prjibelski, A. D. et al. Accurate isoform discovery with IsoQuant using long reads. Nat Biotechnol 41, 915–918 (2023).

6. Tang, A. D. et al. Full-length transcript characterization of SF3B1 mutation in chronic lymphocytic leukemia reveals downregulation of retained introns. Nat Commun 11, 1438 (2020).

7. Chen, Y. et al. Context-aware transcript quantification from long-read RNA-seq data with Bambu. Nat Methods 20, 1187–1195 (2023).

8. Wyman, D. et al. A technology-agnostic long-read analysis pipeline for transcriptome discovery and quantification. Preprint at 10.1101/672931 (2020).

9. Volden, R. et al. Identifying and quantifying isoforms from accurate full-length transcriptome sequencing reads with Mandalorion. Genome Biol. 24, 167 (2023).

10. Kuo, R. I. et al. Illuminating the dark side of the human transcriptome with long read transcript sequencing. BMC Genomics 21, 751 (2020).

11. Pardo-Palacios, F. J. et al. Systematic assessment of long-read RNA-seq methods for transcript identification and quantification. Nat. Methods 21, 1349–1363 (2024).

12. Chen, Y. et al. A systematic benchmark of Nanopore long-read RNA sequencing for transcript-level analysis in human cell lines. Nat. Methods 22, 801–812 (2025).

13. Su, Y. et al. Comprehensive assessment of mRNA isoform detection methods for long-read sequencing data. Nat. Commun. 15, 3972 (2024).

14. Dong, X., et al. The long and the short of it: unlocking nanopore long-read RNA sequencing data with short-read differential expression analysis tools. NAR Genom. Bioinform. 3, lqab028 (2021).

15. Dong, X. et al. Benchmarking long-read RNA-sequencing analysis tools using in silico mixtures. Nat. Methods 20, 1810–1821 (2023).

16. Baid, G. et al. DeepConsensus improves the accuracy of sequences with a gap-aware sequence transformer. Nat. Biotechnol. 41, 232–238 (2023).

17. Hall, M. B. et al. Benchmarking reveals superiority of deep learning variant callers on bacterial nanopore sequence data. Elife 13, (2024).

18. Zolotarov, G. et al. MicroRNAs are deeply linked to the emergence of the complex octopus brain. Sci. Adv. 8, eadd9938 (2022).

19. Zhang, R. et al. A high-resolution single-molecule sequencing-based Arabidopsis transcriptome using novel methods of Iso-seq analysis. Genome Biol. 23, 149 (2022).

20. Zimmermann, B. et al. Topological structures and syntenic conservation in sea anemone genomes. Nat. Commun. 14, 8270 (2023).

21. Occean, J. R. et al. Gene body DNA hydroxymethylation restricts the magnitude of transcriptional changes during aging. Nat. Commun. 15, 6357 (2024).

22. Nejo, T. et al. Challenges in the discovery of tumor-specific alternative splicing-derived cell-surface antigens in glioma. Sci. Rep. 14, 6362 (2024).

23. Leshkowitz, D. et al. Exploring differential exon usage via short- and long-read RNA sequencing strategies. Open Biol. 12, 220206 (2022).

24. Patowary, A. et al. Developmental isoform diversity in the human neocortex informs neuropsychiatric risk mechanisms. Science 384, eadh7688 (2024).

25. Chen, Y. et al. Gene Fusion Detection and Characterization in Long-Read Cancer Transcriptome Sequencing Data with FusionSeeker. Cancer Res. 83, 28–33 (2023).

26. Wright, D. J. et al. Long read sequencing reveals novel isoforms and insights into splicing regulation during cell state changes. BMC Genomics 23, 42 (2022).

27. Zhang, Z., Bae, B., Cuddleston, W. H. & Miura, P. Coordination of alternative splicing and alternative polyadenylation revealed by targeted long read sequencing. Nat. Commun. 14, 5506 (2023).

28. Reese, F. et al. The ENCODE4 long-read RNA-seq collection reveals distinct classes of transcript structure diversity. Preprint at 10.1101/2023.05.15.540865 (2023).

29. Pardo-Palacios, F. J. et al. SQANTI3: curation of long-read transcriptomes for accurate identification of known and novel isoforms. Nat. Methods 21, 793–797 (2024).

30. Paul, L. et al. SIRVs: Spike-In RNA Variants as External Isoform Controls in RNA-Sequencing. Preprint at 10.1101/080747 (2016).

31. Keil, N., Monzó, C., McIntyre, L. & Conesa, A. Quality assessment of long read data in multisample lrRNA-seq experiments using SQANTI-reads. Genome Res. 35, 987–998 (2025).

32. Liu, T. et al. Transcriptome Universal Single-isoform COntrol: A Framework for Evaluating Transcriptome reconstruction Quality. Preprint at 10.1101/2025.08.23.671926 (2025).

33. Pertea, G. & Pertea, M. GFF utilities: GffRead and GffCompare. F1000Res. 9, 304 (2020).

34. Di Tommaso, P. et al. Nextflow enables reproducible computational workflows. Nat. Biotechnol. 35, 316–319 (2017).

35. Barnett, D. W., Garrison, E. K., Quinlan, A. R., Strömberg, M. P. & Marth, G. T. BamTools: a C++ API and toolkit for analyzing and managing BAM files. Bioinformatics 27, 1691–1692 (2011).

36. Goldfarb, T. et al. NCBI RefSeq: reference sequence standards through 25 years of curation and annotation. Nucleic Acids Res. 53, D243–D257 (2025).

37. Li, H. Minimap2: pairwise alignment for nucleotide sequences. Bioinformatics 34, 3094–3100 (2018).

38. Dobin, A. et al. STAR: ultrafast universal RNA-seq aligner. Bioinformatics 29, 15–21 (2013).

39. Iso-Seq - Scalable De Novo Isoform Discovery from Single-Molecule PacBio Reads. (Pacific Biosciences, 2023).

40. Volden, R. et al. Improving nanopore read accuracy with the R2C2 method enables the sequencing of highly multiplexed full-length single-cell cDNA. Proceedings of the National Academy of Sciences 115, 9726–9731 (2018).

